# Replenishing Age-Related Decline of IRAK-M Expression in Retinal Pigment Epithelium Attenuates Outer Retinal Degeneration

**DOI:** 10.1101/2023.09.27.559733

**Authors:** Jian Liu, David A. Copland, Alison J. Clare, Mathias Gorski, Burt T. Richards, Louis Scott, Sofia Theodoropoulou, Ursula Greferath, Katherine Cox, Oliver H. Bell, Kepeng Ou, Jenna Le Brun Powell, Jiahui Wu, Luis Martinez Robles, Yingxin Li, Lindsay B. Nicholson, Peter J. Coffey, Erica L. Fletcher, Robyn Guymer, Monte J. Radeke, Iris M. Heid, Gregory S. Hageman, Ying Kai Chan, Andrew D. Dick

## Abstract

Unchecked, chronic inflammation is a constitutive component of age-related diseases, including age-related macular degeneration (AMD). Here we identified interleukin-1 receptor-associated kinase (IRAK)-M as a key immunoregulator in retinal pigment epithelium (RPE) that declines with age. Rare genetic variants of IRAK-M increased the likelihood of AMD. IRAK-M expression in RPE declined with age or oxidative stress and was further reduced in AMD. IRAK-M-deficient mice exhibited increased incidence of outer retinal degeneration at earlier ages, which was further exacerbated by oxidative stressors. The absence of IRAK-M disrupted RPE cell homeostasis, including compromised mitochondrial function, cellular senescence, and aberrant cytokine production. IRAK-M overexpression protected RPE cells against oxidative or immune stressors. Subretinal delivery of AAV-expressing IRAK-M rescued light-induced outer retinal degeneration in wild-type mice and attenuated age-related spontaneous retinal degeneration in IRAK-M-deficient mice. Our data support that replenishment of IRAK-M expression may redress dysregulated pro-inflammatory processes in AMD, thereby treating degeneration.

**One Sentence Summary:** IRAK-M is a protective molecule and promising therapeutic target for macular degeneration

## INTRODUCTION

Cell-autonomous responses such as metabolic regulation, autophagy and immune-mediated inflammation initiated by noxious stress (environmental factors) are active processes that help to maintain homeostasis (*1*). However, loss of immune regulation and persistent inflammation beget divergent or excessive immune responses, leading to detrimental acute or chronic tissue damage. Such chronic inflammation is accentuated by age (inflammageing) and implicated in progression of many age-related degenerative disorders (*2*).

The retinal pigment epithelium (RPE) is essential for maintenance of outer retinal function and ocular immune privilege. Dysfunction of the RPE leads to photoreceptor (PR) loss and gradual loss of the central visual acuity, as observed in age-related macular degeneration (AMD) (*3–5*). AMD is a progressive, multifactorial disease that is a leading cause of irreversible severe vision loss in the elderly. Alongside ageing, the interplay of oxidative stress and chronic inflammation, resulting from genotype-predisposed susceptibility and environmental stressors, is a significant driver of AMD. Multiple genome-wide association studies have identified risk loci for late and/or early AMD, including, but not exclusively, genes in the complement pathway and *ARMS2/HTRA1* alleles (*6, 7*). Specifically, rare coding variants in regulatory genes of complement such as *CFH* and *CFI* have been associated with AMD risk. This knowledge has led to developing therapeutics, including complement inhibitors and gene therapies for augmenting regulators of complement pathway (*8, 9*). Notwithstanding there remain a number of pathological pathways implicated in the pathogenesis of AMD, including oxidative stress and innate immune responses (*10, 11*). In mice, for example, a high fat diet is required to illuminate pathology on the background of complement gene mutation (*12*). Therefore, elucidating factors central to the diverse pathologies in AMD is critical, irrespective of genetic risk.

Multiple inflammatory pathways are associated with AMD progression, including activation of the complement cascade and NLRP3 inflammasome, production of cytokines and chemokines (e.g., IL-1β, IL-6, IL-8, IL-12, MCP-1, and TNF-α), and low levels of infiltrating cells to the outer retina, such as dendritic cells and macrophages, as well as immune-activated microglia and RPE (*13–17*). Emerging evidence also indicates the association of Toll-like receptors (TLRs), particularly TLR2, 3 and 4, in the risk of development of AMD (*18–20*). The Myddosome is an oligomeric complex consisting of an adaptor protein MyD88 and IL-1R-associated kinase (IRAK) family proteins, and required for transmission of both TLR and inflammasome-IL-1R axis-mediated signals (*16*). Conversely, Myddosome signalling also promotes inflammasome activation (*21*). Although an overactivation of the Myddosome has been observed in the RPE from patients with geographic atrophy (GA, late stage of atrophic AMD) (*16*), important questions remain to be determined such as whether the overactivation has a pivotal role in AMD progression and which component(s) of the Myddosome complex lead to the dysregulation of TLR/IL-1R pro-inflammatory signaling cascades.

Highlighting a central role in the pathophysiology of the retina, the RPE exhibits the highest number of differentially expressed genes (DEGs) overlapping with the genes associated with ageing and age-related retinal diseases and is highly susceptible to the perturbance of ageing and inflammatory stressors (*22*). When the disturbance in RPE intracellular processes, such as autophagy, phagolysosome, mitochondrial metabolism, protein trafficking and senescence, is compounded by oxidative stress, inflammation is elaborated by inflammasome activation and IL- 1β/IL-18 release (*17, 23, 24*). Associated with tissue and organ damage in clinical scenarios such as neurodegeneration, cancer and pulmonary diseases, the magnitude of oxidative stress-induced inflammation is largely determined by various TLRs and balanced by counteracting mechanisms regulated by inhibitors including IRAK-M (gene symbol *IRAK3*) (*25, 26*). Acting as a pseudokinase, IRAK-M downregulates the pro-inflammatory cascade by impeding the uncoupling of phospho-IRAK1/4 from the Myddosome for TGF-β-activated kinase 1 (TAK1)-dependent NF- κB activation, or by forming an IRAK-M/MyD88 complex that stimulates the second wave of NF- κB activation to induce inhibitory modulators (*27, 28*).

IRAK-M is expressed in organs including the liver, heart, brain, spleen, kidney, and thymus (*29*). Downregulation of IRAK-M signalling is associated with exaggerated oxidative stress and systemic inflammation in metabolic disorders such as insulin resistance and obesity. Reduced IRAK-M expression in monocytes and adipose tissues of obese subjects leads to elevated mitochondrial stress, systemic inflammation, and metabolic syndrome (*30*). Multiple mutations in *IRAK3* have been associated with early-onset chronic asthma in humans (*31*).

Following our finding of IRAK-M protein expression in the RPE, a study of IRAK-M in retinal ageing and degeneration was undertaken. We determined the role of IRAK-M in the development of AMD by evaluating genetic variants and their association with AMD risk, evaluating expression of IRAK-M in patient samples and mouse models, and also evaluating changes in retinal function in transgenic mice lacking IRAK-M. Overall, the expression of IRAK-M within human and mouse retinas showed an RPE-specific decline with ageing and was associated with the induction of oxidative stress. RNA-Seq data mining and histology studies divulged a lower IRAK-M expression level in AMD eyes compared to age-matched controls. *Irak3^-/-^*mice developed earlier pathological changes in the retina and RPE with age than wild-type mice, which was accentuated by oxidative stressors. Finally, by overexpressing IRAK-M, we demonstrate a protective role of IRAK-M maintaining RPE cell function and homeostasis, thereby curbing retina degeneration in mouse models.

## RESULTS

### Rare protein-altering variants of *IRAK3* are associated with increased risk of late AMD

In view of the observation of Myddosome activation in AMD (*16*), we asked whether changes in the Myddosome components contribute to disease risk and pathogenic pathways. Analysis of rare variants that alter peptide sequences (non-synonymous), truncate proteins (premature stop), or affect RNA splicing (splice site) can help to identify causal mechanisms – particularly when multiple associated variants reside in the same gene (*32*). Based upon the genetic data from the International AMD Genomics Consortium (IAMDGC) that contains 16,144 late AMD cases versus 17,832 age-matched controls (*6*), we found no genetic association between rare variants of *MYD88* and late AMD (P = 0.95). We then examined the cumulative effect of rare protein-altering variants for all IRAK family kinases (*IRAK1-4*). Among these 4 closely related candidate genes, our analysis highlighted a statistically significant late AMD risk-increasing signal for *IRAK3* (P = 0.012) (Table 1). Table S1 lists the variants in the *IRAK3* gene region, including 18 polymorphic variants that were detected in both AMD cases and controls and used in the gene burden test. As a comparator for *IRAK3*, rare variants of *IL33* that encodes a Th2-oriented cytokine linked to retinal pathophysiology (*33–35*) were not associated with late AMD (P = 0.18).

**Table 1.** Rare protein-altering variants of IRAK3 is associated with increasing risk of late AMD by IAMDGC genomic analysis.

| Gene Symbol | CHROM | Start | End | N Markers | Increasing/decreasing risk for AMD | P value |
| --- | --- | --- | --- | --- | --- | --- |
| *IRAK3* | 12 | 66,582,994 | 66,648,402 | 18 | Increasing risk | 0.012 |
| *IRAK1* | X | 153,278,500 | 153,284,192 | 2 | Increasing risk | 0.35 |
| *IRAK2* | 3 | 10,219,555 | 10,280,654 | 12 | Increasing risk | 0.91 |
| *IRAK4* | 12 | 44,172,041 | 44,177,510 | 3 | Decreasing risk | 0.22 |

The cumulative effect of rare protein-altering variants in the 16,144 late AMD cases versus 17,832 controls of four IRAK genes in the IAMDGC data was examined using gene burden test.

### IRAK-M expression in RPE is reduced with age and to a greater extent in AMD patients

IRAK-M expression was originally reported to be expressed solely by monocytes and macrophages (*36*). As we observed *IRAK3*’s association with late AMD risk, we explored IRAK-M expression in the retina by performing immunohistochemistry on frozen human retinal sections from a young donor eye (20y old, no recorded eye diseases). The data showed an abundant IRAK-M distribution at the RPE layer of the retinal sections, which were co-stained with anti-RPE65 (Fig. 1A) and anti-rhodopsin (Fig. 1B), respectively. Weaker immunopositivity of IRAK-M was found within other retinal layers, including GCL (ganglion cell layer), IPL (inner plexiform layer), OPL (outer plexiform layer), ONL (outer nuclear layer), POS (PR outer segment) and choroid (Fig. 1A and B). Negative controls with primary antibody omitted did not show any signal. An independent immunohistochemistry experiment also demonstrated IRAK-M expression by human RPE in the RPE/choroidal sections from a 73y-old male donor (without recorded eye diseases; Fig. S1A and B). Similar to human expression, IRAK-M was expressed in the mouse RPE (Fig. S1C-E) and a human RPE cell line ARPE-19 (Fig. S1F). These findings are consistent with our previously reported detection of IRAK-M transcript in a murine RPE cell line *in vitro* (*23*). We also observed strong immunopositivity of IRAK-M in both inner (non-pigmented) and outer (pigmented) ciliary epithelium of human eyes (Fig. S1G and H), emphasizing a potential regulatory role in barrier cells.

**Fig. 1.**
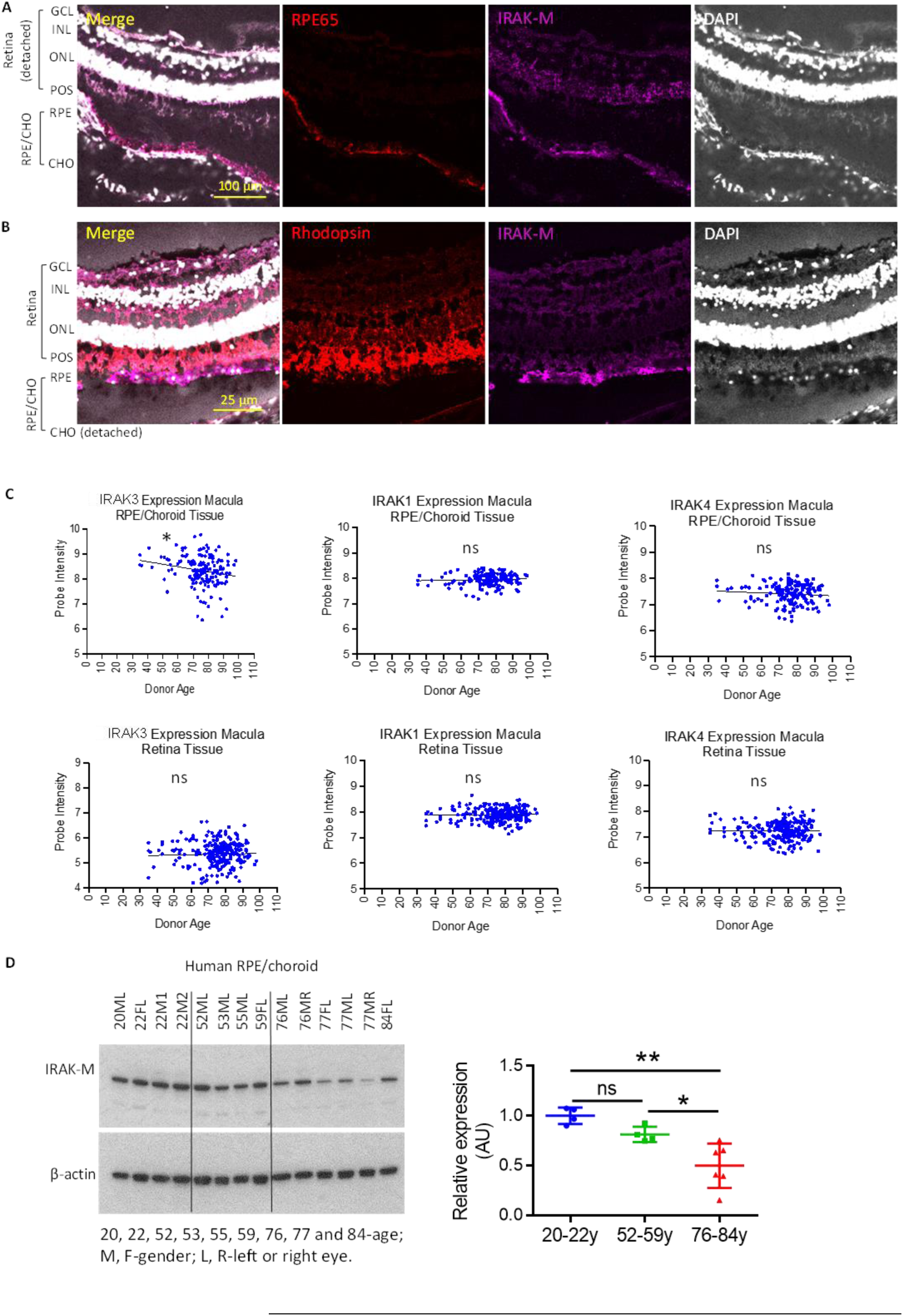
IRAK-M is expressed in RPE and its expression level is reduced with age and in AMD. (**A&B**) Confocal images of human retinal sections from a 20-year-old donor (without recorded ocular disease) demonstrate IRAK-M immunopositivity at the RPE layer (anti-RPE65 stain). DAPI and anti-Rhodopsin were used to stain nuclei and POS, respectively. (**C**) Affymetrix chip-based transcriptome analyses show an age-related reduction in the expression level of *IRAK3* mRNA in macular RPE/choroid tissues, but not in the retina. Neither *IRAK1* nor *IRAK4* mRNA level is changed with age in RPE/choroid or retina. (**D**) Western blot and densitometry quantification show reduced levels of IRAK-M protein expression in aged human RPE/choroidal lysates. The IRAK-M levels were normalized to β-actin (n=4-6). *P < 0.05; **P < 0.01; ns, nonsignificant. Comparison by simple linear regression (C) or one-way ANOVA (D).

We next determined whether the expression level of IRAK-M altered during ageing, the essential pre-requisite for developing AMD. Microarray of human eye samples (without recorded eye diseases) identified an age-dependent decrease in *IRAK3* transcript levels in the macular (Fig. 1C) and extramacular (Fig. S2) RPE/choroid. There was no change in expression in the retina (Fig. 1C and Fig. S2). Neither *IRAK1* nor *IRAK4* altered with age in RPE/choroid or retina (Fig. 1C and Fig. S2). Further analyses of IRAK-M protein expression in human RPE/choroid lysates across a range of ages revealed significant reduction in elderly samples (76-84y) compared to young (20- 22y) and middle-aged (52-59y) samples (Fig 1D). In parallel with reduced IRAK-M protein expression, increased levels of phospho-IRAK4 and NF-κB p65 were detected (Fig. S3A), supporting activation of inflammatory signaling pathways. CFH and C3 protein expression did not change with age (Fig. S3A). As with human samples, RPE isolated from aged mice (19-24m; correlating to a human age of approximately 75 years (*31*)) had lower IRAK-M protein levels compared with younger mice (2-5m, Fig. S3B). The expression of IRAK-M protein in mouse retinal CD11b+ cells (MACS-isolated-microglia and perivascular macrophages) was also reduced with age (Fig. S3C).

We further sought to ascertain whether IRAK-M expression was compromised in AMD, as compared to age-matched controls. We analyzed a published RNA-Seq dataset (GSE99248), which included PORT-normalized counts for both sense and antisense transcripts (*37*). When assessing all *IRAK* family genes, we found that only the level of *IRAK3* mRNA in RPE/choroid/sclera, and not in the retina, was significantly lower in AMD than age-matched controls (Fig. 2A). *IRAK1*, *IRAK2* and *IRAK4* expression, as well as antisense RNAs specific to any *IRAK*, were unchanged between AMD and controls (Fig. 2A). From the same dataset, we also examined the expression of other known genes for negative regulation of TLR/IL- 1R/MyD88/IRAK1/4 signalling (Fig. S4), including *PIN1* (peptidylprolyl cis/trans isomerase, NIMA-interacting 1, which inhibits TLR transcription factor IRF3), *IL1RN* (IL-1R antagonist), *SOCS1* (suppressor of cytokine signaling 1, which induces MAL ubiquitination required for MyD88 activation), *TOLLIP* (Toll-interacting protein, which binds to IRAK1 to induce translocation of TLRs to endosome for degradation), *FADD* (Fas-associated death domain, which interacts with IRAK1/MyD88 to attenuate the signaling), and *PTPN6* (Tyrosine-protein phosphatase non-receptor type 6, which inhibits SYK activation and blocks MyD88 phosphorylation). None of these genes showed any significant difference between AMD and controls.

**Fig. 2.**
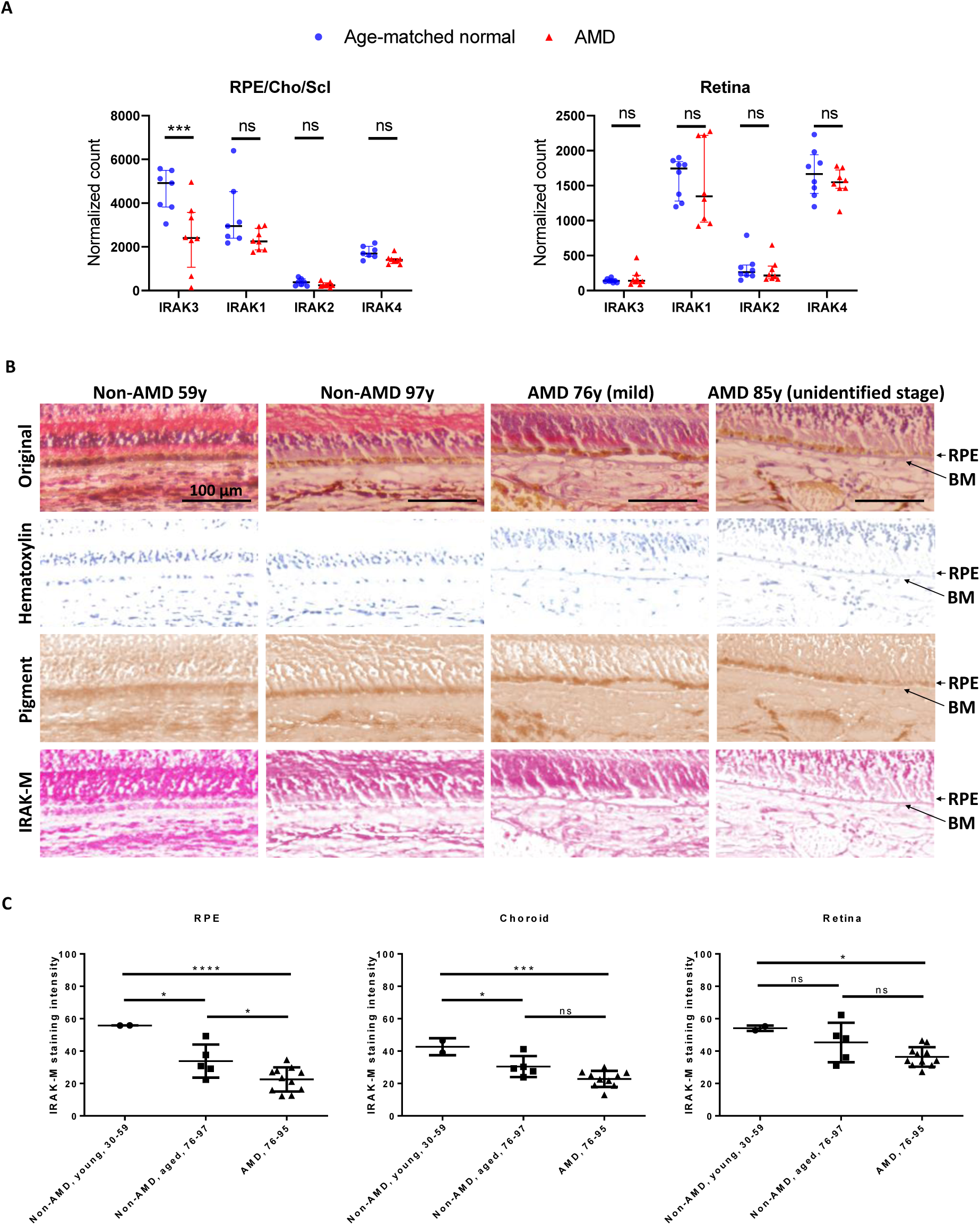
IRAK-M expression level in RPE is reduced in AMD. (**A**) PORT-normalized gene counts from RNA-Seq data (GSE99248) show decreased *IRAK3* mRNA expression in RPE/Choroid/Sclera of AMD donors versus age-matched normal controls. *IRAK1*, *IRAK2* and *IRAK4* mRNA levels in RPE/Choroid/Sclera have no difference. Nor mRNA levels of any *IRAKs* in retina show difference (n=7- 8). (**B**) Magnification of boxed regions of representative IHC images (Fig. S5) of human retinal sections from two non-AMD (59-year and 97-year old, respectively), a mild AMD (76-year old), and an unidentified stage AMD donors (85-year old) were color-deconvoluted using ImageJ to separate IRAK-M staining (red), pigment (brown) and nuclei (blue). Note a nonspecific staining of thickened BM in AMD. (**C**) Quantification of mean staining intensity of macular area shows more severely reduced IRAK-M expression in both aged and AMD RPE, while the reduced expression in choroid is only significant with old age. There are no changes in retina with ageing or in AMD (n=2 for young control, n=5 for old control and n=11 for AMD). *P < 0.05; ***P < 0.001; ****P < 0.0001; ns, nonsignificant. Comparison by two-way ANOVA (A) or one-way ANOVA (C).

To further determine any spatial expression of IRAK-M protein within tissue associated with age and AMD, we performed IHC on paraffin-embedded retinal sections of 2 ‘young’ (aged 30 and 59y) and 5 ‘aged’ (76-97y) individuals without history of AMD, and 11 AMD patients (76-95y). The paraffin slides were visualized using AP-based IHC due to strong autofluorescence of the RPE that was not fully blocked by Sudan black B quenching. In young samples, IRAK-M (stained in red) was observed in various layers of the retina, RPE and choroid (Non-AMD 59y, Fig. S5A and Fig. 2B). In aged control and AMD samples, the pattern and strength of IRAK-M-immunopositive signals was variable, for example with a heightened signal in OPL/ONL (Non-AMD 97y, Fig. S6B and Fig. 2B), in INL/ONL/IS (inner segment) (Mild AMD 76y, Fig. S5C and Fig. 2B), or in NFL (nerve fiber layer) (Unidentified stage of AMD 85y, Fig. S5D and Fig. 2B). After color deconvolution using Fiji package of ImageJ, the IRAK-M signal (red) and RPE pigment (brown) could be separated and discerned for quantification. Advancing our data in Fig. 1C and D), we identified a marked reduction in IRAK-M expression at the macular RPE and choroid with older age (Fig. 2C). The IRAK-M level of expression was also lower in AMD-macular RPE areas compared to age-matched subjects and was not observed in choroid underlying the macula (Fig. 2B and C). Reduction of IRAK-M expression in extramacular tissues was only evident in aged versus young choroid (Fig. S6). Nonspecific staining of Bruch’s membrane (BM) for IRAK-M was observed in AMD samples (Fig. 2B) and in negative staining controls. The intensified BM was not evident in non-AMD eyes (*38*).

### IRAK-M-deficient mice acquire earlier outer retinal degeneration during ageing

Having established the association between reduced IRAK-M expression and age/AMD, we used IRAK-M-deficient mice (without *Rd8* mutation) to investigate whether ageing and lack of IRAK-M affected outer retinal degeneration. The *Irak3^-/-^* mouse line bears an IRAK-M mutant, where two-thirds of the pseudokinase domain (exons 9-11) were removed by homologous recombination (*36, 39*). The multiple conserved cysteine residues within the dimeric structure of the pseudokinase domain of native IRAK-M are essential in the forming of an interactive interface with IRAK4 for the negative regulation of IRAK-Myddosome signaling (*40*).

Pathological changes were tracked for 15 months using fundoscopy and OCT. Between 2 and 5m of age, there was a sharp increase in the incidence of retinas displaying variable number of fundus white spots, from 22.7% (5 out of 22 eyes) to 50% (15 out of 30) (Fig. S7A, Fig. 3A and B). The fundus spots in mice have been well described as a feature of retinal inflammation linked to accumulated macrophages/microglia in the subretinal space (*41, 42*). The incidence of abnormal retinal appearance increased and reached 78.6% of eyes (11 out of 14) by 15m (Fig. 3B). Repeated imaging of the same affected retinas showed that the white spots developed with ageing (Fig. 3A). In comparison, WT mice maintained normal retinal appearance (i.e., no progression of white spots) at 12m, however a substantial incidence of WT retinas displayed white spots between 12 and 21m as mice aged (61.5% or 16 out of 26 at 19-21m, Fig. 3B). These time course data demonstrate accelerated ageing-associated retinal abnormalities and degeneration associated with defective IRAK-M (Fig. 3B). Notably, the early appearance of retinal spots was accompanied by outer retinal lesions identified by OCT (Fig. 3C).

**Fig. 3.**
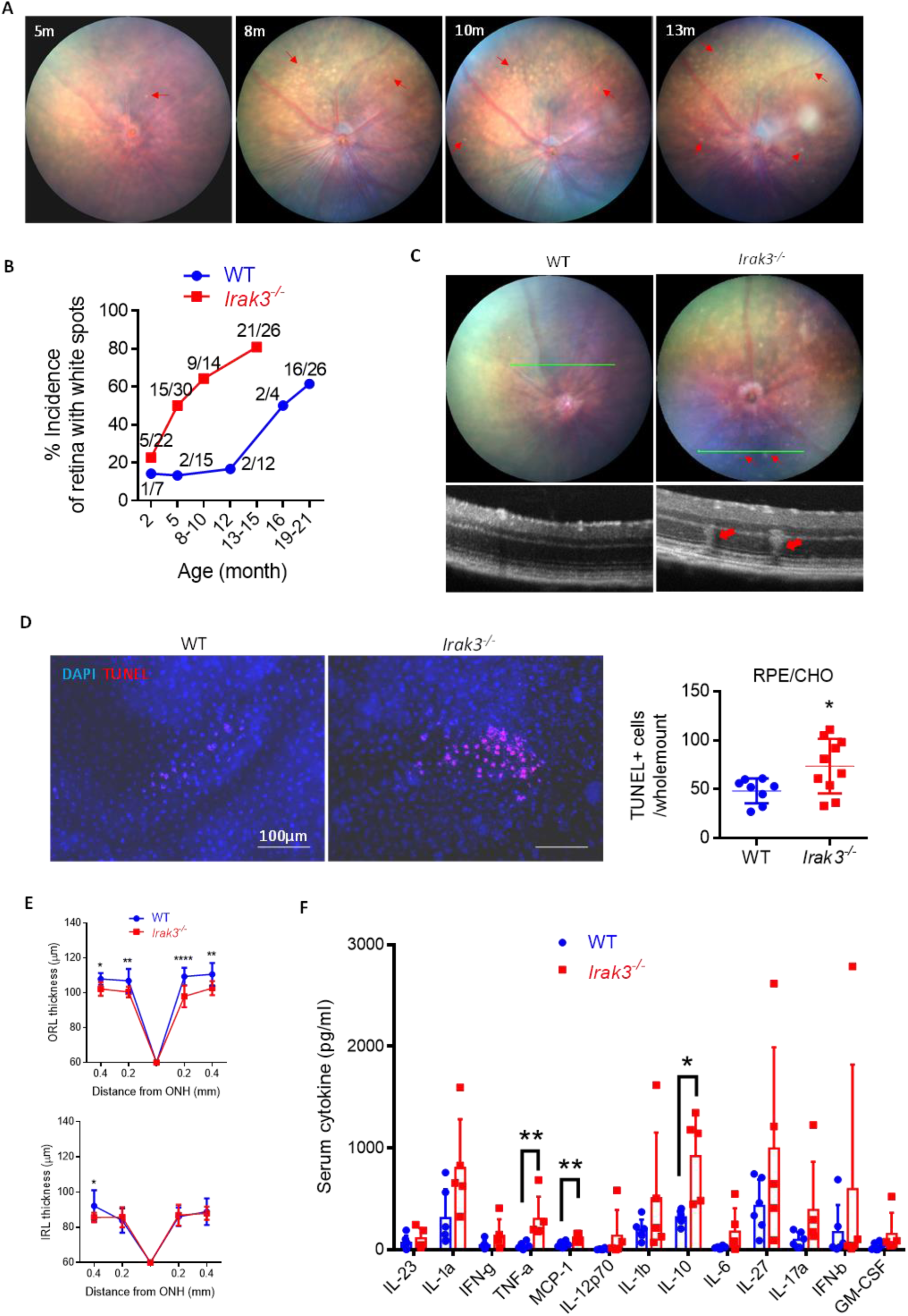
*Irak3^-/-^* mice spontaneously display early retinal abnormalities. (**A**) Representative fundal images show age-related appearance of white spots (red line arrow) in *Irak3^-/-^* mouse retinas. (**B**) Time course of incidence of flecked retina (number of spots > 3) shows increased incidence of retinal spots in *Irak3^-/-^* mice compared to WT controls. Each value is a ratio of number of flecked retina to total number of retina at each time point. (**C**) Representative fundal and OCT images demonstrate that the white spots (red line arrow) are associated with outer retinal abnormalities (red arrow) in 5m-old *Irak3^-/-^* mice. (**D**) TUNEL staining on RPE/choroidal flatmounts reveals elevated number of apoptotic cells in *Irak3^-/-^* mice versus WT controls (5m-old) (n=8-10). (**E**) Quantification of OCT images indicates significant outer retinal thinning in *Irak3^-/-^* mice aged 12-13m. The change in inner retinal thickness is negligible (n=6-12). (**F**) Multiplex cytokine array demonstrates an overall higher levels of serum cytokines in *Irak3^-/-^* compared to WT mice (12-13m- old), where the increases of TNF-α, MCP-1 and IL-10 serum concentrations are statistically significant (n=5-6). *P < 0.05; **P < 0.01; ****P < 0.0001. Comparison by unpaired two-tailed Student’s t-test (D) or two-way ANOVA (E and F).

Increased numbers of CD11b+ myeloid cell populations in the outer nuclear layer (ONL) (Fig. S7B), and CD11b+ cell accumulation in the subretinal space (Fig. S7C) were observed in *Irak3^-/-^* mice, associated with increased number of apoptotic cells (TUNEL-positive) within the RPE/choroid (Fig. 3D). Although no difference in retinal thickness was found at 5m between WT and *Irak3^-/-^* mice, the outer retina of *Irak3^-/-^*mice was significantly thinner by 12-13m (Fig. 3E). In parallel, by 12-13m serum inflammatory cytokine levels in *Irak3^-/-^* mice were higher than in the WT mice (significant increases in TNF-α, MCP-1 and IL-10; Fig. 3F).

### Oxidative stress reduces RPE-IRAK-M expression and loss of IRAK-M increases susceptibility of outer retina to oxidative damage

Age-associated accumulation of oxidative stress in the RPE is a recognised contributor to the progression of AMD. To examine if oxidative stress could be an independent factor for the reduction of IRAK-M expression, we applied oxidative stressors both *in vitro* and *in vivo*.

*In vitro*, a human ARPE-19 cell line was treated with different doses of paraquat (PQ), a stable chemical primarily inducing mitochondrial ROS, for up to 72h. LDH cytotoxicity assay showed a dose-dependent cytotoxicity caused by PQ exposure for 72h (Fig. S8A), whereas IRAK-M protein expression was suppressed by a sub-toxic dose of PQ (0.25mM) (Fig. S8B). Reduction in IRAK-M was accompanied by an enhanced pro-inflammatory response, demonstrated by the increased secretion of pro-inflammatory cytokines HMGB1, IL-18 and GM-CSF, and decreased secretion of anti-inflammatory IL-11 (Fig. S8C). Likewise, downregulation of IRAK-M expression level following 72h treatment of sub-toxic doses of PQ (0.25-0.5mM) occurred in human iPSC-derived RPE (Fig. S8D-F) and human primary RPE cells (Fig. S8G and H).

*In vivo*, retinal oxidative damage was introduced by fundus camera-directed light exposure (100kLux for 20min) (*43*) or intravitreal administration of PQ (2µl at 1.5mM) (*44*) in C57BL/6J WT mice aged 8w. Western blot analyses showed that IRAK-M expression in the RPE lysate was significantly abated after 7 days in both models (Fig. 4A and D). Fundoscopy and OCT photographs obtained on day 14 displayed the fundal appearance of white spots (red arrows, Fig. 4B and E) indicative of accumulated microglia/macrophages inside the ONL (*42*), alongside thinning of the outer retina indicative of cell loss in the light-induced retinal degeneration (LIRD) model (Fig. 4C), and reduced thickness in both outer and inner retina in the PQ model (Fig. 4F).

**Fig. 4.**
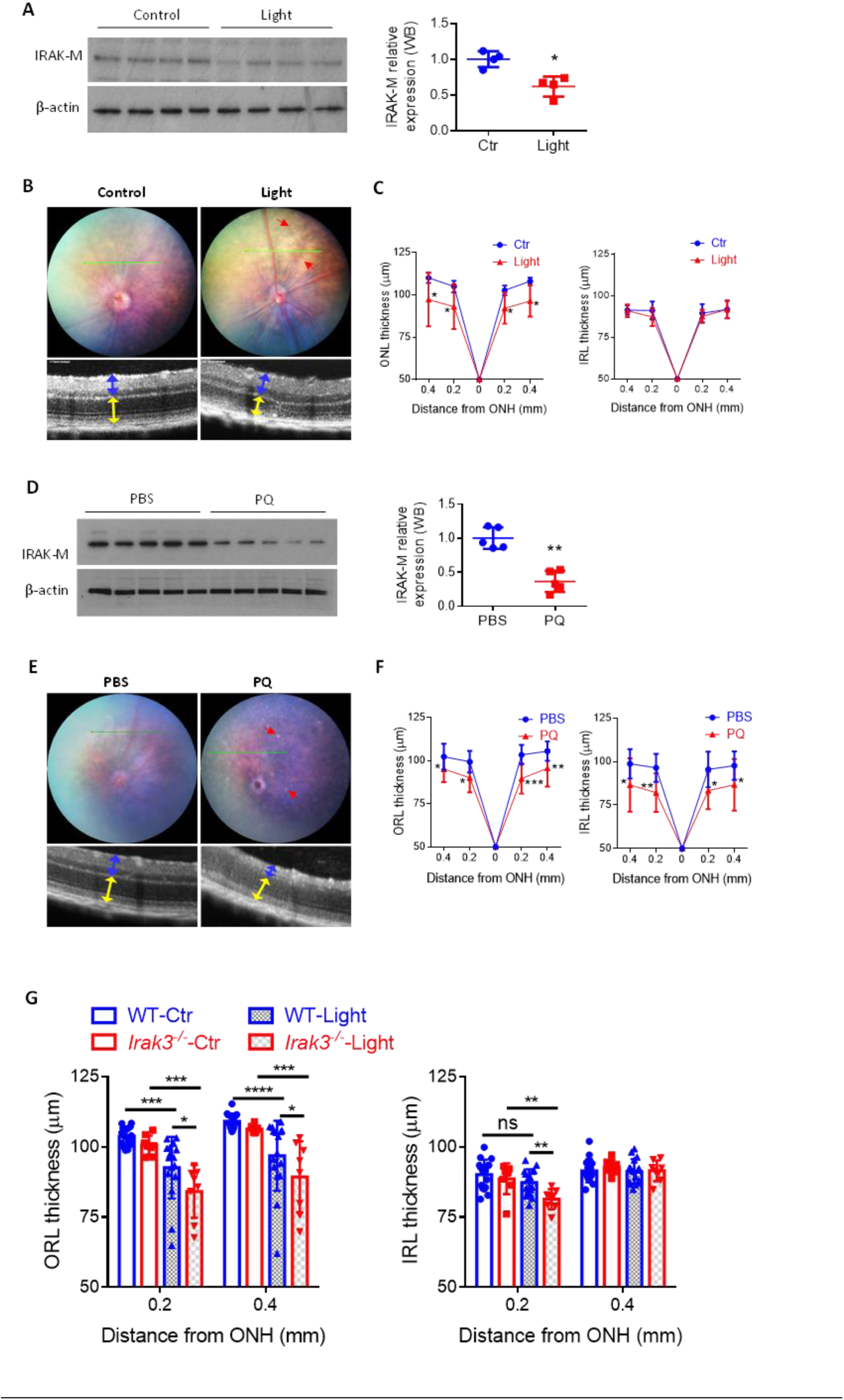
Wild-type mice exhibit reduced RPE-IRAK-M expression level by oxidative stress and *Irak3^-/-^* mice are more vulnerable to light-induced retinal degeneration. Retinal oxidative stresses were induced in 8-week-old C57BL/6J mice by either fundus-light induction (100kLux for 20min, **A-C**) or intravitreal administration of paraquat (PQ, 2µl at 1.5mM, **D-F**). (**A&D**) Western blot analyses of IRAK-M expression in RPE lysate on day 7 post oxidative damage (n=4 or 5). (**B&E**) Representative fundoscopy and OCT images obtained on day 14 demonstrate appearance of retinal lesions (red line arrows), and reduced thickness of outer retina (yellow double-arrow lines) in light model (n=8, **C**), or both outer and inner retina (blue double-arrow lines) in PQ model (n=9-11, **F**). (**G**) Eight-week-old WT and *Irak3^-/-^* mice were subjected to retina oxidative insults by light induction. OCT quantification of retinal thickness (average of temporal and nasal measurements) demonstrates exaggerated retinal thinning in *Irak3^-/-^* mice compared to WT controls on 14 days post light induction, which is more pronounced in outer retinal layers (n=8-16). *P < 0.05; **P < 0.01; ***P < 0.001; ****P < 0.0001. Comparison by unpaired two-tailed Student’s t-test (A and D) or two-way ANOVA (C, F and G).

Given the observed age-dependent increase of retinal pathology in IRAK-M-depleted mice, we next explored whether oxidative stress would exaggerate the effect. Retinal oxidative stress was induced in adult WT and *Irak3 ^-/-^* mice (8w old) by light induction. *Irak3^-/-^* mice exhibited amplified retinal damage compared to WT, particularly a thinner outer retinal layers following light challenge (Fig. 4G).

### AP-1 regulates IRAK-M expression in RPE cells in age-dependent manner

Known transcription factors regulating IRAK-M expression in monocytes or lung epithelial cells include activation protein 1 (AP-1) and CCAAT/enhancer-binding protein beta (C/EBP-β) (*45, 46*). By analysing human RPE/choroidal lysate derived from donor eyes without recorded ocular disease, we found that along with age-associated reduction in IRAK-M level, expression of c-Jun, an AP-1 subunit, was decreased in aged samples compared to young controls (Fig. S9A and B). c-Fos, another AP-1 subunit, was reduced in old age compared to middle-age samples (Fig. S9A and B). C/EBP-β expression had no change during ageing process.

The association of c-Jun and c-Fos with the IRAK-M promoter region were confirmed by ChIP assay on ARPE-19 cells, and this was significantly enhanced in response to LPS stimulation for 24h (Fig. S9C). To investigate whether oxidative stress altered AP-1 activity or expression in the RPE, we treated ARPE-19 cells with PQ and demonstrated a dose-dependent downregulation of phosphorylation of both c-Jun and c-Fos after 72h, while total c-Jun and c-Fos expression were downregulated by higher dose of PQ (Fig. S9D and E). Through inhibition of AP-1 subunit activity and expression, SP600125 (primarily targeting c-Jun) and T5224 (targeting c-Fos) at 20 µM significantly decreased IRAK-M expression (Fig. S9D and E). Consequently, treatment with AP- 1 inhibitors resulted in enhanced ARPE-19 susceptibility to PQ-induced cytotoxicity (Fig. S9F), similar to the observed effect induced by IRAK-M siRNA (Fig. S9G). Increasing c-Jun expression via CRISPR/Cas9 activation plasmid upregulated IRAK-M expression (Fig. S9H). The effect on suppressing oxidative stress-induced cytotoxicity by overexpressing IRAK-M using CRISPR/Cas9 activation plasmid transfection was not found when c-Jun expression was augmented (Fig. S9H).

### IRAK-M deficiency induces RPE mitochondrial dysfunction and senescent phenotype which is protected by IRAK-M augmentation

To elucidate metabolic mechanisms involved in IRAK-M deficiency-induced retinal degeneration, we examined RPE cell metabolism and senescence using primary mouse RPE cells. IRAK-M-deficient cells showed reduced levels of basal mitochondrial respiration (BR) and ATP production compared to WT cells as assessed by OCR analyses (Fig. S10A), while no significant differences in basal glycolysis (BG) and maximal glycolytic capacity (MGC) were observed between genotypes as assessed by ECAR (Fig. S10B). These data infer a role of IRAK-M in the maintenance of mitochondrial function in RPE cells. In support of this, *Irak3 ^-/-^*RPE cells were more prone to oxidative stressor (PQ or H_2_O_2_)-induced senescent phenotype, marked by increased SA-β-gal activity (Fig. S10C), enhanced expression of cyclin-dependent kinase inhibitor p21^CIP1^, decreased nuclear lamina protein LB1 (Fig. S10D), and elicited secretion of IL-6 (a senescence-associated cytokine) (*17*) (Fig. S10E). The basal secretion level of pro-inflammatory cytokine HMGB1 of *Irak3^-/-^* RPE cells was significantly higher than the WT cells but the responsiveness to the oxidative stressors were comparable (Fig. S10F).

Based upon the data above demonstrating a role of IRAK-M in the context of ageing and oxidative challenge, we examined whether an overexpression of IRAK-M could protect RPE. Native IRAK-M expression in human iPSC-derived RPE cells was augmented via transfection of a CRISPR/Cas9-based activation plasmid (Fig. S11A). After 48h of transfection, the cells were treated with H_2_O_2_ or LPS for a further 24h. OCR analysis demonstrated that basal and maximal mitochondrial respiration were both sustained by IRAK-M overexpression, but impaired in sham-transfected cells following oxidative or immune stresses (Fig. 5A). Although untreated IRAK-M-overexpressing iPSC-RPE cells displayed lower maximal glycolytic activity than control plasmid-transfected cells, the level remained stable upon H_2_O_2_ or LPS treatment (Fig. 5B). In contrast, glycolytic activity in control cells was significantly reduced by 24h treatment with H_2_O_2_ or LPS (Fig. 5B). The lower level of glycolysis in un-stressed iPSC-RPE with overexpressed IRAK-M suggests less bio-energetic dependency on glucose, with possible benefits to glucose-dependent photoreceptors (*47*).

**Fig. 5.**
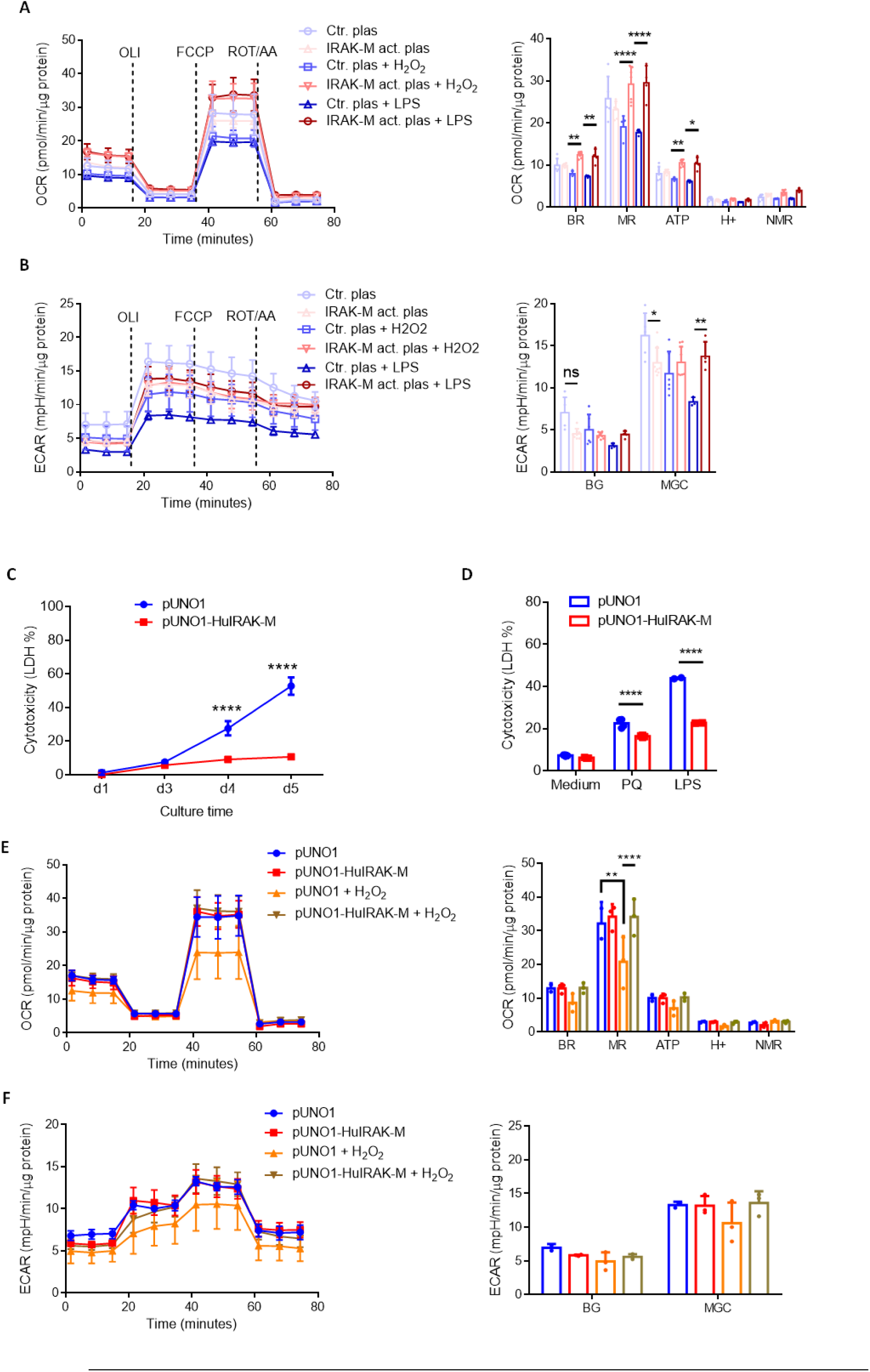
Overexpression of IRAK-M in RPE cells supports metabolic activities and inhibits cell death against stressors. (**A&B**) Metabolic flux analyses demonstrate that increasing endogenous IRAK-M expression in human iPSC-RPE cells via CRISPR/Cas9 activation plasmid maintains both mitochondrial respiration (OCR, **A**) and glycolytic capacity (ECAR, **B**), upon 24h treatment with 30 µM H2O2 or 1 µg/ml LPS (n=3-7). (**C**) Stably transfected cell lines selected from mouse B6-RPE07 cells were established to persistently express human IRAK-M. Time course of LDH release over 5 days since confluence of monolayers shows sustained cell viability by human IRAK-M transfection (n=4-8). (**D**) Human IRAK-M expression inhibits PQ (125 µM) or LPS (40 ng/ml)-induced cytotoxicity post 72h of treatment in stably transfected B6-RPE07 cells (n=2-4). (**E&F**) Primary mouse *Irak3^-/-^*RPE cells were subjected to transient transfection for human IRAK-M expression using pUNO1 plasmid and 48h later, the cells were treated with 60 µM H2O2 for another 24h. OCR analysis (**E**) shows protected mitochondrial maximal respiration by human IRAK-M against oxidative stress treatment. ECAR analysis (**F**) does not show any changes in glycolysis activity by H2O2 treatment or IRAK-M transfection (n=3). *P < 0.05; **P < 0.01; ****P < 0.0001; ns, nonsignificant. Comparison by two-way ANOVA.

ARPE-19 cells with IRAK-M overexpression induced by CRISPR/Cas9 partially reversed LPS-induced reduction in maximal mitochondrial respiration (Fig. S11B and C), supporting the findings from human iPSC-RPE cells (Fig. 5A). Taken further, overexpression of IRAK-M in ARPE-19 promoted the formation of autophagosomes (LC3B-GFP) and autolysosomes (LC3B- RFP) following H_2_O_2_ or LPS treatment, suggesting an upregulated autophagy flux (Fig. S11D). Moreover, ARPE-19 senescence induced by sub-toxic dose PQ (0.25 mM) was prevented by IRAK-M overexpression, as we documented decreased SA-β-gal activity and HMGB1 secretion (Fig. S11E and F). Finally, a marked LDH release induced by a toxic dose of PQ (1 mM) was significantly subdued by increasing IRAK-M expression (Fig. S11G).

We then created stably transfected RPE cell lines maintained in selective medium from a parent mouse B6-RPE07 cell line that expressed either mouse or human *IRAK3* mRNA (Fig. S12A). Expression of mouse *Irak1* and *Irak4* were not affected. A NF-κB activity assay showed a decrease in DNA-binding activity of nuclear NF-κB in human *IRAK3*-expressing mouse cells after LPS stimulation (Fig. S12B), demonstrating that the transduced human *IRAK3* is as functional as its murine counterpart in suppressing NF-κB activation in mouse RPE. Stably transfected RPE cells overexpressing human *IRAK3* survived longer, compared to sham-transfected cells when assessing cell death after four days of confluency (Fig. 5C). Freshly confluent cells (with stable overexpression of *IRAK3*) exhibited a reduced stressor-induced cytotoxicity after treatment with PQ (0.125 mM) or LPS (40 ng/ml) for 3 days (Fig. 5D). To exclude the possible contribution of native mouse *Irak3* to cell response observed, we performed transient transfection on primary RPE cells isolated from *Irak3^-/-^*mice. A metabolic flux assay was applied to examine metabolic alterations in response to shorter period of treatment with H_2_O_2_ (24h, Fig. 5E and F). Similar to data from human iPSC-RPE cells using CRISPR/Cas9 activation plasmid (Fig. 5A and B), maximal mitochondrial respiration in mouse primary *Irak3^-/-^* RPE cells was retained by human *IRAK3* transduction after H_2_O_2_ treatment (Fig. 5E). H_2_O_2_ -induced oxidative stress had no effect on glycolysis in *Irak3^-/-^* RPE cells (Fig. 5F).

### AAV2-mediated IRAK-M expression suppresses light-induced retinal degeneration in wild-type mice and spontaneous retinal degeneration in *Irak3^-/-^* mice

To correct defective gene expression or function in diseases, experimental approaches have included introducing human genes, such as *RPE65*, *CFH* and *ND4* (NADH dehydrogenase subunit 4), to mouse eyes for functional or preclinical evaluation (*48–51*). Such studies have utilised AAV2 and translated to clinical trials to treat RPE-related eye diseases (*52*). To identify the dose-dependent transduction efficacy, 2 µl of AAV2 encoding EGFP under the control of constitutive cytomegalovirus (CMV) promoter (AAV2.CMV.EGFP) at 1×10^12^ or 2×10^11^ gc/ml were delivered into mouse eyes via the subretinal route. The ‘high dose’ (1×10^12^ gc/ml in 2 µl, or 2×10^9^ gc/eye) induced a more pronounced EGFP expression 2-11 weeks post the injection than the “low dose” (2×10^11^ gc/ml or 4×10^8^ gc/eye) (Fig. S13A). Administration with AAV2.CMV.hIRAK3 induced a dose-dependent *IRAK3* mRNA expression in RPE/choroid two weeks post injection, compared to a similar vector but with no transgene used as a control ‘null’ vector (Fig. 6A). Transduced human IRAK-M protein was detected in the RPE, as demonstrated by immunohistochemistry, using two independent IRAK-M antibodies (Fig. 6B).

**Fig. 6.**
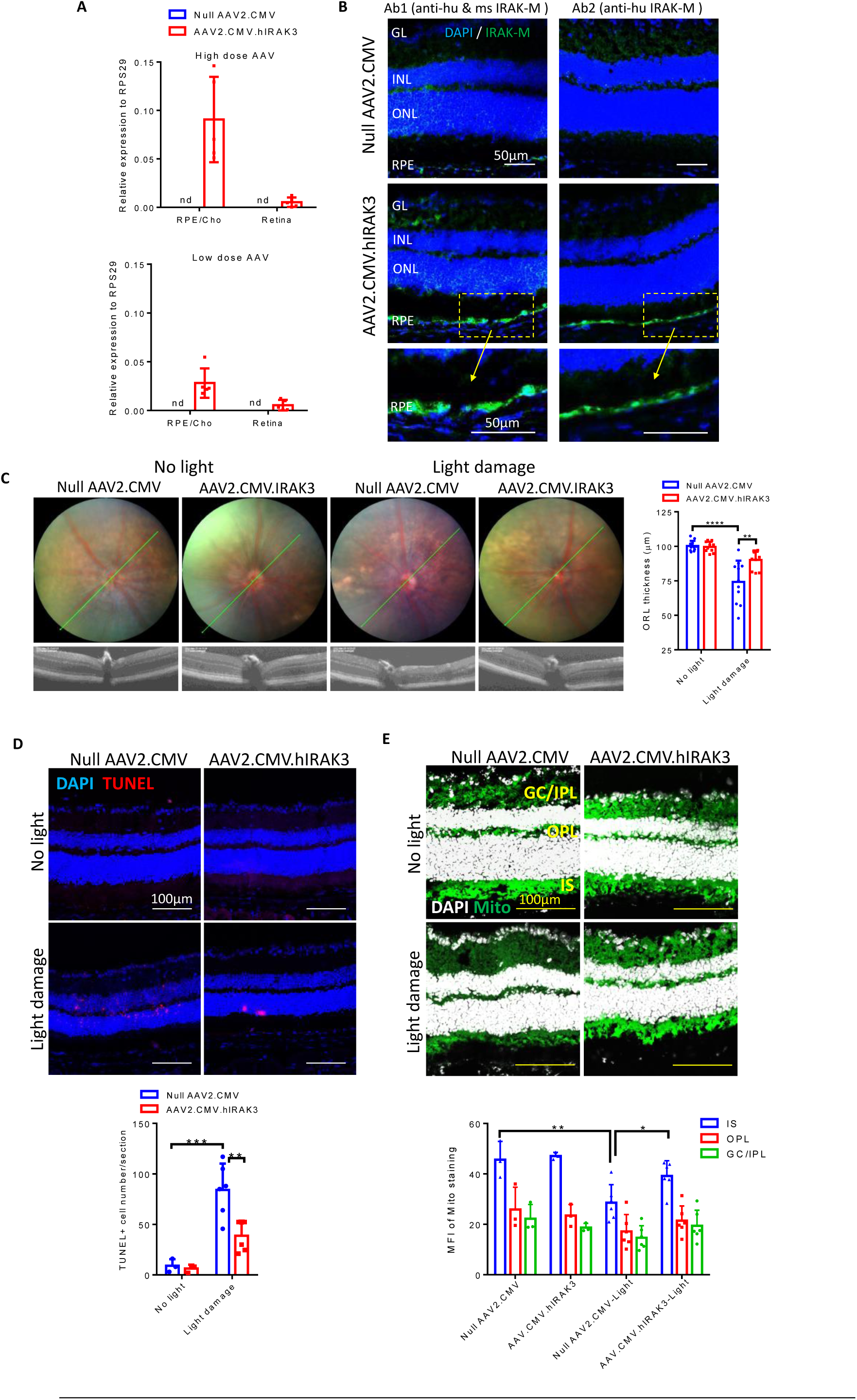
Subretinal delivery of AAV.hIRAK3 protects retina against light damage in wild-type mice. (**A**) Two weeks post subretinal injection of AAV2.CMV.hIRAK3 or AAV2.CMV (high dose 2×10^9^ versus low dose 4×10^8^ gc/eye), RPE/choroid and retina were analyzed for IRAK3 transgene expression using qRT- PCR, normalized by RPS29 mRNA (n=5). (**B**) Retinal cryosections were examined for high dose AAV- mediated IRAK-M expression using an antibody recognizing both human and mouse IRAK-M (Ab1), or an antibody specific to human IRAK-M (Ab2). Representative confocal images were shown. (**C-E**) Two weeks after subretinal injection with the high dose of AAV2.CMV.hIRAK3 or null vector, each mouse was subjected to light-induced retinal degeneration in one eye and left thereafter for a further two weeks, followed by assessment of retinal pathology and therapeutic response. (**C**) Representative fundoscopy/OCT images and quantification show light-induced retinal lesions and averaged outer retinal thickness (n=10- 11). (**D**) Representative confocal images of TUNEL staining on retinal sections and quantification of 3 sections from each eye (n=3-6). (**E**) Confocal images of MitoView Green staining for mitochondrial content and MFI measurement in 3 different fields from two sections of each eye (n=3-6). *P < 0.05; **P < 0.01; ***P < 0.001; ****P < 0.0001. Comparison by two-way ANOVA.

To evaluate the protective effects of IRAK-M transgene expression *in vivo*, we applied light- induced retinal degeneration in mice 2 weeks after AAV injection (2×10^9^ gc/eye). Light exposure of the null AAV2-injected eyes resulted in a decrease of outer retinal thickness, indicative of the PR loss. The protective effect of AAV2.CMV.hIRAK3 treatment from PR injury was conspicuous, as demonstrated by suppression of light-induced outer retinal thinning (Fig. 6C). The LIRD model exhibited significant outer retinal thinning (Fig. 4C), supported by TUNEL+ apoptosis in the ONL (Fig. 6D). There were fewer number of TUNEL+ cells within inner retinal layers, indicating a secondary cell death response in the inner retina following PR loss (*53, 54*). Contemporaneous with the retaining of retinal thickness by AAV.IRAK3 was a reduction in light-induced TUNEL+ cell apoptosis within retinal sections (Fig. 6D), as well as an inhibition of mitochondrial impairment in the IS (Fig. 6E). The mitochondria in GL, IPL and OPL were less affected by light challenge (Fig. 6E).

Based upon our finding that *Irak3^-/-^ mice* developed signs of retinal degeneration earlier than WT, we asked whether AAV-IRAK3 could attenuate outer retinal degeneration caused by IRAK-M deficiency and ageing. To this end, we performed subretinal administration of AAV2.CMV.hIRAK3 or null AAV2.CMV (2×10^9^ gc/eye) in young *Irak3 ^-/-^* mice (2-4m old) and allowed them to age. Six months following the subretinal delivery of AAV vectors, we found that AAV2.CMV.hIRAK3 blunted the age-dependent occurrence of retinal spots (Fig. 7A and B), and significantly reduced the number of retinal spots in aged *Irak3 ^-/-^* mice compared to the null vector (8-10m old; Fig. 7C). The effect was more pronounced within the treatment side of the retina receiving the vector, as expected (Fig. 7C). Importantly, compared to the null AAV treated mice, AAV-IRAK3 delivery showed attenuated outer retinal thinning (Fig. 7D).

**Fig. 7.**
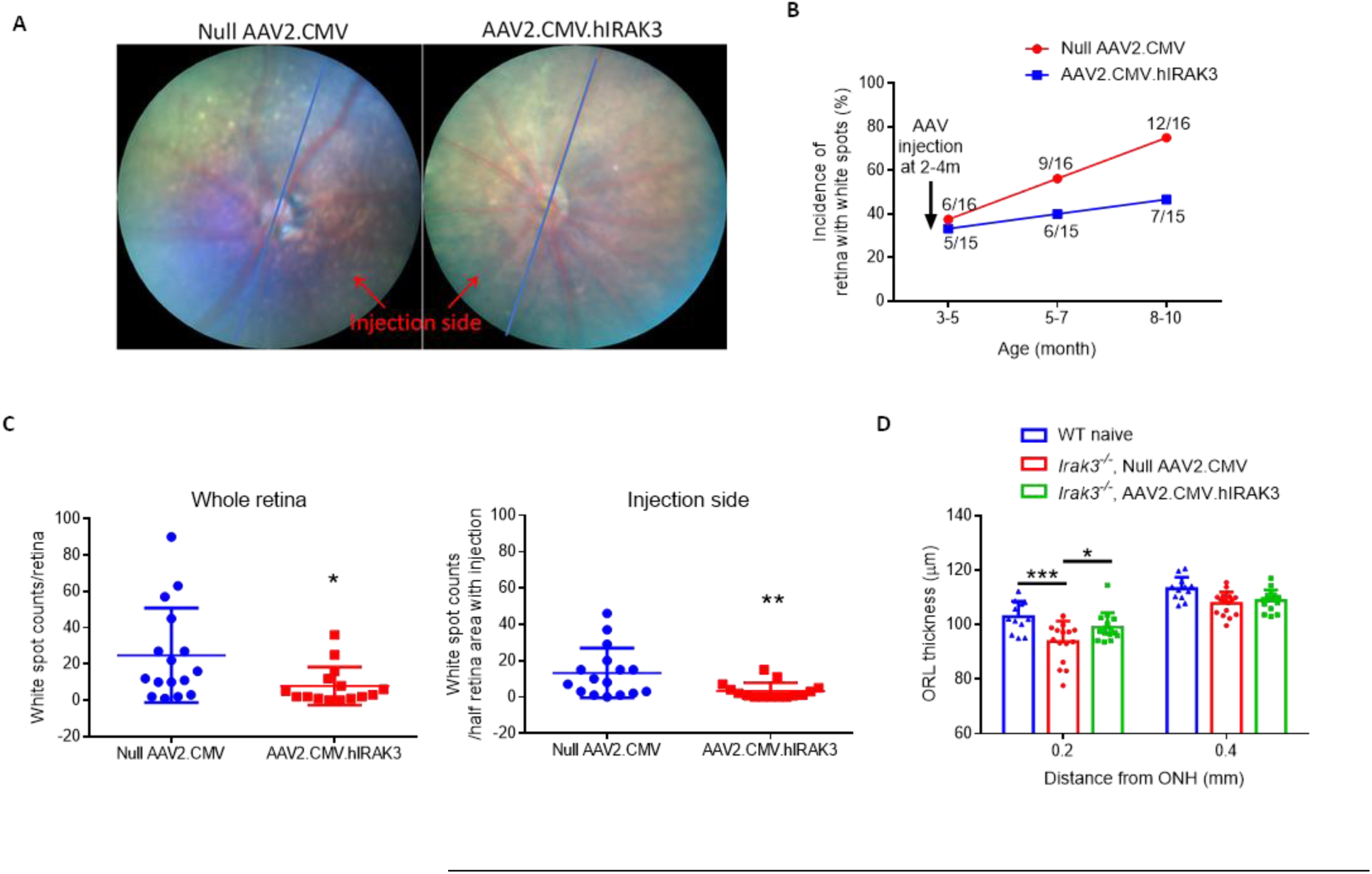
Subretinal delivery of AAV.hIRAK3 prevents from age-related spontaneous retinal degeneration in *Irak3^-/-^* mice. 2×10^9^ gc of AAV2.CMV.hIRAK3 was injected subretinally in one eye of each *Irak3^-/-^* mouse (2-4m old), with null vector injected to the contralateral eye. Mice were then monitored by fundoscopy and OCT for 6 months thereafter. (**A**) Representative fundal images show retinal spots in 8m-old *Irak3^-/-^* mice with AAV administration at the age of 2m. Blue lines separate the retina into two sides based on the injection site. (**B**) Time course of incidence of flecked retina shows IRAK3 gene therapy decelerated the appearance of retinal spots in ageing *Irak3^-/-^* mice (n=15 or 16). (**C**) Number of retinal spots in whole retina or at the injection side, was blind-counted for comparison between AAV2.CMV.hIRAK3 and null vector groups (8-10m-old, n=15 or 16)). (**D**) OCT quantification shows a reduction in outer retinal thickness close to the centre region (0.2 mm distant from optic nerve head) in 8-10m-old *Irak3^-/-^*mice compared to age-matched WT littermates, which is revoked by AAV.hIRAK3 gene delivery (n=12-16). *P < 0.05; **P < 0.01; ***P < 0.001. Comparison by unpaired two-tailed Student’s t-test (C) or two-way ANOVA (D).

## DISCUSSION

Among the plethora of pathways implicated in AMD, there is a strong association and evidence base for a central role of altered immune responses and innate immune dysregulation alongside pro-degenerative stressors, such as oxidative stress and metabolic perturbation. In the present study, we have demonstrated a protective role of the immune regulator IRAK-M in the metabolic and immune homeostasis of the RPE. This is based on the expression levels of IRAK-M in young, old and AMD human eyes, genetic variant burden in those with AMD and experimental models of oxidative stress, ageing and IRAK-M deficiency. A feed-forward loop with ageing, oxidative stress and expression decline of the immune regulator, IRAK-M, may constitute a pro-inflammatory microenvironment driving retinal degeneration. Replenishing the homeostatic regulator IRAK-M maintains mitochondrial function, inhibits pro-inflammatory senescence and promotes cell survival, therefore protecting the retina from degeneration in a LIRD model and progressive degeneration in *Irak3^-/-^* mice. As we observed, IRAK-M is consistently reduced with ageing, oxidative stress and AMD; the replenishment of IRAK-M may be a broadly applicable therapeutic strategy for treating AMD patients.

Prior work has noted the expression of IRAK-M in cells other than monocytes/macrophages, including airway and intestine epithelium, fibroblasts, neurons, neutrophils, dendritic cells basophils and B cells (*36, 55, 56*). In lung biopsy samples from healthy humans, IRAK-M is highly expressed in type II epithelial cells and the dysfunction of IRAK-M is implicated in inflammatory lung diseases (*31*). Tarallo *et al.* reported aberrant activation of NLRP3-inflammasome and Myddosome signaling, such as increased phospho-IRAK1/4 levels in RPE lysates of GA patients, albeit without probing the regulator IRAK-M (*16*). Here we report that the expression of IRAK-M declines with age in the RPE but not retinal tissue and is reduced further in AMD subjects compared to age-matched controls (Fig. 1, 2, S2, S3, S4 and S6). Additionally, we found that IRAK-M was expressed by bilayer ciliary epithelium (Fig. S1G and H), indicating the distribution of this key inflammation inhibitor in other ocular epithelium barriers. The RPE regulates and protects against excessive oxidative stressors, inflammasome activation, mitochondrial impairment, lipid accumulation and cellular senescence (*4, 17, 26, 57*), all pathways that can propel the insidious AMD progression (*13, 14*). TLRs (TLR1-7, 9 and 10) are expressed by RPE cells and IL-1Rs are ubiquitously distributed (*58*). Coupled with the known immunosuppressive factors produced by the RPE, such as membrane molecules CD200, IL-1R2, IL-1Ra, FasL, and anti-inflammatory chemokines or cytokines (CX3CL1, TGF-β, IL-11 and IFN-β) (*23, 59–62*), IRAK-M is required for balancing the regional innate and adaptive immune activation and suppression at the posterior segment of the eye. We showed that *Irak3^-/-^*mice incurred greater oxidative damage, including RPE cell mitochondrial dysfunction, pro-inflammatory senescence, and early AMD-like pathologies such as subretinal accumulation of myeloid cells, outer retinal lesions, and cell death (Fig. 3, 4G, S7 and S10). Additionally, *Irak3^-/-^*mice displayed systemic inflammation evidenced by increased serum cytokine levels.

The downregulation or upregulation of IRAK-M expression is context-dependent. For instance, upregulation of IRAK-M was identified following ischemia-reperfusion of liver and brain (*63, 64*), and in infarcted heart (*65*), where it is thought to limit the magnitude of immune responses and repair pro-inflammatory damage. In a mouse model of cerebral ischemia, IRAK-M was found to be induced by HIF1α and played a neuroprotective role by inhibition of NF-κB signaling and production of COX-2, TNF-α, NLRP3 and iNOS. In comparison, *IRAK3^-/-^* mice developed exacerbated infarcts (58). In contrast to acute responses, downregulation of IRAK-M was more associated with chronic diseases, exemplified by alcoholic liver disease, inflammatory bowel disease, insulin resistance and metabolic syndrome (*25, 29, 30*). Indeed, whilst acute alcohol intake increases IRAK-M expression in human monocytes, chronic alcohol exposure results in its decrease in expression and enhanced inflammation (*66*). In obese subjects, reduced IRAK-M levels in monocytes and adipose tissues constitute a causative factor of mitochondrial oxidative stress and systemic inflammation (*30*). Furthermore, age-related decreases in the basal level of IRAK-M and its inducibility upon TLR activation have been discovered in PBMCs and fibroblasts in rodents (*67, 68*). Using RNA-Seq, Western blot and IHC, we localized the decline in IRAK-M expression to the RPE, rather than of the retina or choroid, in ageing, oxidative stress and AMD (Fig. 1C-D, 2A-C, 4A-B, S2, S3B), indicating that RPE-IRAK-M serves as an early harbinger molecule of degeneration progressing to AMD. Increasing IRAK-M in the RPE *via* boosting endogenous gene expression or exogenous gene delivery helped to maintain cell functions (mitochondrial activity and autophagy) and inhibit cellular senescence and NF-κB activity (Fig. 5, S11 and S12), implying the importance of IRAK-M for the RPE health (Fig. 6 and 7).

Reduced IRAK-M expression with age may be a repercussion of the pathophysiological processes in the genomic or epigenomic programmes. Our data show an association of reduced AP-1 subunit proteins c-Jun and c-Fos and decreased IRAK-M expression (Fig. S9). This agrees with the findings that aged human fibroblasts display declined c-Jun and c-Fos proteins, a shifted distribution of AP-1 components and DNA binding capacity (*69*). Decreased transcription activity of AP-1 has been linked to tissue and cell ageing (*70, 71*), as opposed to NF-κB which was frequently increased in activity in aged tissues and age-related illnesses such as Alzheimer’s disease, diabetes and osteoporosis (*71, 72*). A recent genome-wide profiling study indicated that AP-1 functioned as a governing agent in the senescence programme by shaping the enhancer landscape and determining the dynamic hierarchy of the transcription factor network leading to senescence (*73*). In our work, AP-1 inhibition in human ARPE-19 cells and IRAK-M-deficient murine primary RPE cells rendered the cells more susceptible to oxidative stress-induced cytotoxicity and/or senescence. Of note, and possibly due to the miscellaneous functions of AP- 1/c-Jun signalling in stress response and apoptosis (*74, 75*), increasing c-Jun expression or activity had no beneficial effect on oxidative damage protection (Fig. S9H).

Limitations exist in this study and further investigations will enable a deeper mechanistic understanding of retinal degeneration and help inform potential therapeutic approaches. Our *in vivo* assessment did not reveal any ocular toxicity when overexpressing IRAK-M for more than 6 months in *Irak3^-/-^* mice. However, for translation assessment, large animal studies will be required. We overexpressed IRAK-M by different methods, including CRISPR activation and plasmid- or AAV-based gene delivery, and observed benefits. Future studies should determine levels of IRAK-M expression to define the dose-response curve and further interpret the role of IRAK-M regulation and levels of AP-1 activity (*29*).

In conclusion, we have identified an age-related diminishment of IRAK-M expression largely restricted to the RPE, which is worsened in AMD. Our findings offer insights into a previously unrecognized mechanism where IRAK-M plays a crucial role to maintain RPE cell homeostasis and function via co-targeting mitochondrial health, oxidative stress, autophagy and inflammation. As a consequence, gene augmentation of IRAK-M demonstrates translational benefit in counteracting side-effects of ageing or oxidative stress and reducing outer retinal degeneration in disease models. Given the complexity of multiple affected pathways in AMD, a therapeutic strategy via manipulating IRAK-M in the RPE to address multiple pathways is potentially applicable in a wider population of AMD patients.

## MATERIALS AND METHODS

### Study Design

The overall goals of this study were to define whether alteration of IRAK-M expression in RPE during the ageing process and in AMD occurs. The subsequent goal was to develop a targeted gene therapy for age-related and inflammation-driven RPE and retinal degeneration. The primary experimental procedures are described below, with detailed Materials and Methods listed in the Supplementary Materials.

#### Human sample analyses

For investigations on human ocular samples in all respective institutions, experiments were conducted according to the Declaration of Helsinki principles and in compliance with approved institutional guidelines. We used gene burden test on the large-scale genetic data from International AMD Genomics Consortium (IAMDGC) that contains 16,144 late AMD cases versus 17,832 age-matched controls (*6*) to analyze whether there was a genetic association between rare protein-altering variants of IRAK-M and AMD, compared to other Myddosome-associated proteins. Human age-related progressive changes in mRNA expression of IRAKs were probed in samples including 227 extramacular; 159 macula RPE/choroid and 238 extramacular; 242 macula retina 6 mm trephine tissue punches, by Affymetrix chip-based Microarray and linear regression analysis. Age-related change of IRAK-M expression in RPE/choroid at the protein level was determined by Western blot using postmortem eye tissues from young, middle-aged and aged groups with mixed genders (4-6 samples per age group). AMD-associated changes in mRNA expression of IRAKs and known genes involved in the negative regulation of TLR/IL- 1R/MyD88/IRAK1/4 signalling pathways were discerned by data mining of RNA-Seq data (GSE99248), containing 8 AMD donor eyes aged 83-95 years versus 7 control donor eyes aged 83-92 years. AMD-associated IRAK-M protein expression change was examined by immunohistochemistry of postmortem eye sections from 11 individuals with varying stages of AMD pathology (aged 76-95 years, mixed gender) and 5 age-matched control subjects without recorded eye disorders (aged 76-97 years, mixed gender). The processing and staining of all sections were executed at the same time with the same vials of reagents and antibody to avoid batch effects.

#### Irak3^-/-^, ageing mice and oxidative stress induction

We used *Irak3^-/-^* and WT mice to define whether ageing and/or lack of IRAK-M affected outer retinal degeneration. As the original *Irak3^-/-^* breeding pairs purchased from Jackson Laboratory (strain B6.129S1-Irak3tm1Flv/J, stock no. 007016) presented Rd8 mutation of Crb1 gene that was not reported previously, the mice were backcrossed with WT C57BL/6J for selection of Rd8- negative *Irak3^-/-^* genotype (*76*). Only male mice from the established Rd8-negative *Irak3^-/-^* colony were used to avoid possible sex-associated variation in immune responsiveness (*77*). All animal experiments were approved by the University of Bristol Ethical Review Group and conducted in accordance with the approved institutional guidelines. Time course of clinical examinations on retinal pathology, including retinal structure, fundus spots and thickness, was performed using Micron IV-guided fundoscopy and optical coherence tomography (OCT) in *Irak3^-/-^* mice (aged 2- 15 months) and WT mice (aged 2-21 months). Primary endpoints were RPE cell death, subretinal accumulation of macrophages, and serum cytokine concentrations at indicated time points. To determine whether oxidative stress could be an independent factor affecting IRAK-M expression, we applied oxidative stressors to different RPE cells *in vitro* and 8-week-old WT mice *in vivo.* The mice were subject to fundus camera-directed light exposure (100kLux for 20min) (*43*) or intravitreal injection of paraquat (2µl at 1.5mM) (*44*). The contralateral eye was left without light challenge or injected intravitreally with PBS as a control. The sample size was chosen empirically based on the results of previous studies, which varied between experimental settings. In general, 4-30 replicates for each condition were used per time point or experiment, with precise numbers specified in the figure legends.

To elucidate metabolic mechanisms involved in IRAK-M deficiency-induced retinal degeneration, we isolated primary RPE cells from 5-month-old *Irak3^-/-^* versus WT littermates and characterized cell metabolism and senescent phenotype. To demonstrate whether IRAK-M had a protecting role for RPE cells against oxidative or immune challenges *in vitro*, we overexpressed IRAK-M by either endogenous CRISPR activation or exogenous IRAK-M delivery via plasmid vectors. *In vitro* cell responses to stressors and IRAK-M gene delivery were assessed for mitochondrial respiration and glycolytic activities, autophagy flux, cytokine secretion, and expression of senescence markers.

#### Therapeutic approaches

We undertook *in vivo* therapeutic evaluation of IRAK-M replenishment via subretinal administration of AAV2-expressing human IRAK-M in two different murine models of retinal degeneration, light-induced outer retinal degeneration in young WT mice and spontaneous outer retinal degeneration in ageing *Irak3^-/-^* mice. In both models, null AAV2 vehicle injections served as a negative control to determine baseline responses. The control AAV2 and IRAK3-expressing AAV2 were both under the control of constitutive cytomegalovirus (CMV) promoter. A pilot experiment to determine viral dose-dependent transduction efficacy was performed by subretinal injection of 2×10^9^ gc (high dose) or 4×10^8^ gc (low dose) of AAV2.CMV.EGFP to each eye and evaluated by fundal fluorescence imaging for 11 weeks. AAV-mediated human IRAK-M transgene expression in mice RPE/retina was verified by qPCR and immunohistochemistry of retinal samples. For the light model, retinas were exposed to light challenge at two weeks post AAV injection, and retinal pathologies were examined after a further two weeks by fundoscopy, OCT, and histology for TUNEL+ cell death and mitochondrial content. For the *Irak3^-/-^* model, we monitored *Irak3 ^-/-^*mice (2-4m old) for 6 months following subretinal injection of AAV vectors using quantitative parameters such as retinal fundus spots and outer retinal thickness, measured by fundoscopy and OCT. Laterality of injected eyes was randomized, and the investigators were blinded to the vector type throughout intervention and analysis.

### Statistics

Results are presented as means ± standard deviation (SD). A simple linear regression was utilized to analyze the correlation between gene expression and human ageing using Microarray data. Statistical analysis was performed using an unpaired two-tailed Student’s t-test between two groups. Tests for normal distribution and homogeneity of variance and comparisons between more than two groups were conducted using one-way ANOVA. A two-way ANOVA was used to assess the interrelationship of two independent variables on a dependent variable, followed by the Kruskal–Wallis test with Bonferroni correction for *post hoc* comparisons. Differences between groups were considered significant at P < 0.05. Statistical analyses were conducted using GraphPad Prism 8.0.

## Author contributions

Conceptualization: ADD, JL and YKC. Methodology: JL, ADD, YKC, GSH, IMH, MG, BR, UG, MJR, ELF, RG, PJC, DAC and LBN. Investigation: JL, YKC, DAC, AJC, MG, BTR, GSH, LS, ST, UG, KC, OHB, KO, JLBP, JW, LMR and YL. Visualization: ADD, JL and YKC. Supervision: ADD, JL and YKC. Writing—original draft: JL, ADD and YKC. Writing—review & editing: all authors.

## Data and materials availability

All data are available in the main text or the supplementary materials.

Fig. S1 to S13

