## Supplementary materials for "Replenishing Age-Related Decline of IRAK-M Expression in Retinal Pigment Epithelium Attenuates Outer Retinal Degeneration"

### Supplementary Materials and Methods

#### *Gene-based burden analysis*

Single variant analyses have limited power to depict rare variants with association. Gene-based burden tests evaluating accumulated association from multiple rare variants per gene have been shown to complement such analyses and improve power to detect a burden of disease. The burden of disease was computed in the IAMDGC (*1*) using the variable threshold test (*2*) as implemented in EPACTS. These analyses assume that all variants in a gene either increase or decrease disease risk. When variants with opposite directions of effect reside in the same gene, power will be reduced. The analysis was focused on protein-altering variants, since it was assumed that the other (not protein-altering) variants would outnumber these predicted deleterious variants by far and would thus dilute a disease burden from the deleterious variants. Assuming a negative selection against such deleterious variants that cause their frequency to be low across ancestries, variants with MAF < 1% in African, Asian, and European ancestry were analyzed. Genotypes of these rare protein-altering variants were utilized if genotyped directly or rounded imputed allele dosages to the next best genotype if imputed; imputed variants were restricted to those of highest imputation quality (RSQ >= 0.8). Statistical significance was assessed by adaptive permutation testing with variable thresholds (up to 100 million permutations; minimal P-value = $1\times10^{-8}$).

#### *Human ocular tissues and sample processing for protein analyses*

For human ocular tissue experiments conducted in the University of Bristol, human donor eyes or ocular tissue surpluses to corneal transplantation (without recorded ocular disease) were obtained from National Health Service (NHS) Blood and Transplant Services after research ethics committee approval (20/LO/0336), with experiments conducted according to the Declaration of Helsinki and in compliance with UK law. The age and sex of human eye samples are indicated in figures. For immunohistochemistry of frozen sections, the tissue was fixed in 4% formaldehyde in PBS, embedded in Optimum Cutting Temperature (OCT) Compound and frozen in dry ice followed by storage at −80°C before preparation of cryosections (8 µm sections). Slides were stored at −80°C until required. For western blotting, dissected RPE/choroid tissue was crushed in 400 µL Pierce® RIPA buffer containing Protease and Phosphatase Inhibitors (Thermo Fisher Scientific, Paisley, UK) for protein extraction.

For human ocular tissue experiments conducted in the University of Melbourne, all procedures and protocols adhered to the tenets of the Declaration of Helsinki and were approved by the Human Ethics Committee of the Royal Victorian Eye and Ear Hospital, Melbourne, Australia. Written informed consent was obtained from patients after the nature and possible consequences of the study were explained. During surgery, immediately after exenteration, the eyes globe was dissected and the anterior contents removed. The posterior eyecups then were placed into a solution containing 4% paraformaldehyde in 0.1 M phosphate buffer (PB) for 4 hours. The time between exenteration and fixation was approximately 10 minutes. The tissues were protected in graded sucrose solutions (10%, 20%, 30% weight/volume), embedded in OCT Compound and cryosectioned (14 μm sections). Slides were stored at −80°C until required.

#### *Human ocular tissues and sample processing for microarray*

For human ocular tissue experiment conducted at the University of Utah, the study was reviewed and approved by the institutional review board at the University of Utah and conforms to the tenets of the Declaration of Helsinki. Written informed consent was obtained from the surviving relatives of all human eye donors or by the donors prior to time of death. We are grateful to the families of all the eye donors for their generous gifts, without which this work would not have been possible. Total RNA was harvested from 227 extramacular; 159 macula RPE/choroid and 238 extramacular; 242 macula retina 6 mm trephine tissue punches using a PAXgene Tissue RNA kit (Qiagen, Germantown, Maryland, USA) following the manufacturer's protocol. Microarray analyses were performed using a proprietary (Diaxonhit) Affymetrix chip with 1.5 million probe sets (4 unique probes per set) targeting every exon, every exon-intron and exon-exon junction for known and predicted transcripts of all genes based on Human Genome NCBI build 37.3. Total RNA was PCR amplified using a NuGEN Ovation Pico WTA System (Tecan Life Sciences, San Jose, California, USA) and cDNA purified with a QIAquick PCR Purification System (Qiagen). Purified cDNA was then fragmented and labeled with biotin using a NuGEN Encore Biotin Module (Tecan Life Sciences). Hybridization of biotinylated cDNAs to microarray chips was performed using standard microarray procedures. Probe intensities were determined using an Affymetrix GeneChip System 3000Ex v.2 (Thermo Fisher Scientific) and resulting CEL files were imported into Genomics Suite analysis software (Partek). Intensity levels of all probes were corrected for background using RMA background correction and normalized to array averages using the quantile normalization function. Probe intensity levels are reported on a log2 scale with a minimum threshold score of 4.5 required to be considered above background. IRAK3, IRAK1 and IRAK4 probe intensity data were queried using Partek software and plotted using GraphPad 9.0 Prism software.

#### *Data mining*

Potential datasets for analyses of AMD-related changes were chosen according to availability of processed RNA-Seq data via the NCBI GEO Datasets, using search terms 'AMD retina' and 'AMD RPE'. The Kim *et al.* RNA-Seq dataset in retinal and RPE-Choroid-Scleral (RCS) Homo Sapiens tissues (GEO accession GSE99248) was used, which includes PORT-normalized antisense and sense transcriptome counts from 7 donor eyes without recorded ocular diseases (age range 83-92y, 3 females and 5 males), and 8 AMD donor eyes (age range 83-95y, 5 females and 2 males). The AMD eyes were characterized at various stages, including 2 early AMD, 1 dry AMD with RPE atrophy, 3 late dry AMD, and 1 late wet AMD. The dataset samples were sorted according to tissue-type (either retina or RCS) and phenotype (AMD or normal); normal samples corresponding to each tissue type were used as controls.

#### *Mice*

*Irak3*$^{-/-}$ mice were obtained from Jackson Laboratory (B6.129S1-Irak3tm1Flv/J, stock #007016). Not reported previously (*3*), we found Rd8 mutation of Crb1 gene within this strain using an established PCR genotyping protocol (*3*). Therefore, the mice were backcrossed with wild-type (WT) C57BL/6J (Charles River Laboratories, Portishead, UK) and Rd8-negative *Irak3*$^{-/-}$ genotype was confirmed (data not shown). The breeding colonies were maintained in the Animal Services Unit of the University of Bristol. All mice were kept in the animal house facilities of the university according to the Home Office Regulations. Treatment of the animals conformed to the Association for Research in Vision and Ophthalmology (ARVO) statement for the Use of Animals in Ophthalmic and Vision Research. The methods were conducted in accordance with the approved University of Bristol institutional guidelines and all experimental protocols under Home Office Project Licences 30/2745 and PP9783504 were approved by the University of Bristol Ethical Review Group.

#### *Antibodies*

Rabbit polyclonal anti-IRAK-M (Cat. ab8116), rabbit polyclonal anti-c-Fos (phospho T325; ab27793), rabbit monoclonal anti-c-Fos (ab134122), rabbit polyclonal anti-HMGB1 (ab18256), rabbit monoclonal anti-c-Jun (ab32137), rabbit monoclonal anti-c-Jun (phosphor S63; ab32385), rabbit monoclonal anti-p21 (ab109199), rabbit monoclonal anti-Lamin B1 (ab133741), mouse monoclonal anti-Ezrin (ab4069) and mouse monoclonal anti-Rhodopsin (ab98887) were all purchased from Abcam (Cambridge, UK). Mouse monoclonal anti-RPE65 was from Abcam (ab78036) or from Chemicon (MAB5428). A rabbit anti-human IRAK-M (HPA043097) was from Merck (Gillingham, UK). Rabbit polyclonal anti-ZO-1 (#61-7300) was from Thermo Fisher Scientific. Rat monoclonal anti-CD11b (M1/70; #550282) was from BD Biosciences (Wokingham, UK). Secondary antibodies used in western blotting, including HRP-conjugated goat anti-rabbit and anti-mouse IgG were from New England Biolabs (Hitchin, UK). Secondary antibodies used in immunostaining including Alexa Fluor 488-goat anti-rabbit or anti-mouse IgG, Alexa Fluor 488-rabbit anti-rat IgG, Alexa Fluor 555-goat anti-rabbit IgG, and Alexa Fluor 488-donkey anti-goat IgG were from Thermo Fisher Scientific.

#### *RPE cell lines, Primary cells, iPSC derived RPE cells and treatment*

A human RPE cell line ARPE-19 (ATCC) and a mouse RPE cell line B6-RPE07 (a gift from Prof. Heping Xu, Belfast) were maintained in DMEM medium supplemented with 10% FBS, 1% L-glutamine, 1 mM sodium pyruvate, 60 μM 2-mercaptoethanol, 1% penicillin/ streptomycin (complete medium) at 37°C in an atmosphere of 5% $CO_2$. Cells were passaged with a split ratio of 1:5 using trypsin/EDTA and allowed to recover for 2 days in complete medium prior to treatment.

Primary murine RPE cells were isolated and cultured as previously reported (*4*). Briefly, eyes from WT or *Irak3*$^{-/-}$ mice were enucleated and cleaned using angles scissors to ensure no connective tissue remained. After cornea and lens were removed, the eyes were incubated at 37°C in hyaluronidase for 45 min, and in Hank's balanced salt solution (HBSS) with 10 mM HEPES for a further 30 min before retinas removed via incision. Following incubation at 37°C in trypsin/EDTA for 45min, eyecups were then transferred into HBSS with 20% heat-inactivated FBS and shaken gently to allow the RPE to detach. The RPE sheets were further incubated in trypsin/EDTA for 1 min to allow the formation of single cell suspensions. The resulting RPE cells were resuspended in alpha MEM basal medium supplemented with 1% N1 Medium Supplement, 1% L-glutamine, 1% penicillin–streptomycin and 1% nonessential amino acid solution (NEAA), 20 µg/l hydrocortisone, 250 mg/l taurine, 0.013 µg/l triiodo-thyronin and 5% FBS. Cell purity was confirmed by immunoblotting for RPE65 and rhodopsin as we described before (*5*). The cells were seeded to laminin-precoated Seahorse XF cell culture plates (Agilent Technologies, Santa Clara, California, USA) or 8-well chamber slides (Corning GmbH, Wiesbaden, Germany) at a density of 25,000/$cm^2$. After the first 72 h of incubation, serum was removed from the medium. The culture medium was changed twice a week. The cells were used for experiments 7-10 days post isolation.

Primary human RPE cells (H-RPE) were purchased from Lonza (Slough, UK). The cells were maintained in RPE Basal Medium supplemented with 2% L-glutamine, 0.5% FGF-B and 0.1% GA-1000, and sub-cultured at ratio of 1:3 using trypsin/EDTA for no more than 4 passages. The H-RPE cells were plated to 24-well plates at a density of 10,000 cells/$cm^2$. After overnight incubation, the cells were used for induction of oxidative stress.

Human fibroblast-induced pluripotent stem cells (iPSCs) derived from a healthy donor were a kind gift from Prof. Peter Coffey from UCL. iPSC colonies were cultured on Matrigel hESC-qualified matrix (BD Biosciences, Wokingham, UK) in E8 (Thermo Fisher Scientific). Once 80% confluent, the media was changed to Differentiation Media containing Knockout-DMEM, 20% Knockout Serum Replacement, 1% non-essential amino acids (NEAA), 1% Glutamax and 0.2% 2-mercaptoethanol (all from Thermo Fisher Scientific). Cultures were fed twice weekly and for at least a further 8 weeks until pigmented foci were observed. These pigmented foci were isolated manually and seeded to Matrigel-precoated 96-well plates or Seahorse XF plates with a density of 50,000 cells/$cm^2$ for induction of oxidative or immune stress and analyses of metabolic function.

The cells from different sources were either continuously treated with different concentrations of paraquat (PQ), hydrogen peroxide ($H_2O_2$) or lipopolysaccharide (LPS, all from Sigma-Aldrich, Poole, UK) for up to 72h, or under a repeated exposure to PQ or $H_2O_2$ for 2h every day for a total of 7 days (*6*). In some experiments, RPE cells were pretreated with a c-Jun inhibitor (SP600125, Sigma-Aldrich) or c-Fos inhibitor (T-5224, Cambridge Bioscience, Cambridge, UK) for 2h before addition of PQ.

#### *CD11b$^+$ cell isolation by magnetic-activated cell sorting (MACS)*

Microglia and perivascular macrophages were isolated by enriching CD11b$^+$ cells from single-cell suspensions of retina tissues from different ages of WT mice, using anti-CD11b microbeads (Miltenyi Biotec, Surrey, UK) according to the manufacturer's instruction. The cells were lysed in Pierce® RIPA buffer containing Halt™ Protease and Phosphatase Inhibitors (Thermo Fisher Scientific) for Western blot analysis of IRAK-M expression.

***Immunohistochemistry and fluorescence staining***

To examine the expression pattern of IRAK-M in human and mouse frozen sections, human eye tissue from a 20-year-old donor (no recorded ocular disease) or enucleated eyes from 8-week-old WT mice were fixed with 4% (for human tissue) or 2% (for murine tissue) paraformaldehyde (PFA) before 12-μm thick cryosections prepared on a cryostat. The sections were permeabilized with 0.1% Triton X-100, blocked with 10% normal donkey serum, 5% BSA plus 0.3 M glycine before incubation with rabbit anti-IRAK-M (1:1000) and mouse anti-Rhodopsin (1:500) or mouse anti-RPE65 (1:100) overnight at 4°C. After wash, sections were incubated with donkey anti-rabbit IgG conjugated with Alexa Fluor 555 and donkey anti-mouse IgG with Alexa Fluor 647 (both 1:1000). DAPI counterstain was used to show nuclei in sections. Tissues were washed and mounted in Vectashield antifade medium and examined by confocal microscopy.

For immunohistochemistry of human paraffin eye sections, paraffin-embedded human eye sections from AMD and non-AMD subjects were purchased from the Lions Gift of Sight (Minnesota, USA) after research ethics committee approval (20/LO/0336). The slides were deparaffinized and rehydrated, followed by antigen retrieval with citrate buffer (pH 6.0) at 90°C for 20 minutes. After three washes in PBS and blocked and permeabilized in 5% normal goat serum (NGS), 5% BSA and 0.1% Triton X-100, specimens were incubated with rabbit anti-IRAK-M antibody (1:250, Cat. ab8116, Abcam, Cambridge, UK) at 4°C overnight. The secondary antibody, a biotinylated goat anti-rabbit IgG (1:1000, Thermo Fisher Scientific, Paisley, UK) visualized via the avidin-biotin-alkaline phosphatase complex (ABC-AP) method (Vectastain ABC-AP Kit, 2Bscientific, Upper Heyford, UK) using the Vector Red Substrate (2Bscientific). Slides were then counterstained by Hematoxylin, dehydrated, and mounted with Histomount medium. The absence of staining when the primary antibody omitted was used as negative control. IHC images at the macular area and peripheral retina were captured using an Evos XL Core microscope (Thermo Fisher Scientific). The labelling of the sections was performed at the same time and samples were imaged using an EVOS XL Core Imaging System with identical settings. Images were processed using Colour Deconvolution plugin in ImageJ to separate Hematoxylin (blue), IRAK-M (AP-Red) and pigment (brown). The ROI of pigmented RPE was carefully identified in the pigment/brown picture, which was copied-and-pasted into the identical area of the AP-Red picture. The mean staining intensity of IRAK-M for RPE was measured using ImageJ. The ROI for retina or choroid was selected based on the nuclear staining (blue).

To prepare mouse retinal and RPE/choroid wholemounts, enucleated eyes were initially fixed in 2% PFA overnight. After dissection of the eyes, the retinal and RPE/choroidal tissues were blocked and permeabilized in 10% normal goat serum, 5% BSA, 0.3 M glycine with 0.3% Triton X-100 in PBS for 2 hours, followed by incubation with the rat anti-CD11b (1:200) in 1% BSA with 0.15% Triton X-100 at 4°C overnight. After thorough wash, samples were further incubated with Alexa Fluor 488-goat anti-rat IgG (1:400). Tissues were counterstained with DAPI and flat-mounted for observation by confocal microscopy.

Cell apoptosis in the retinal and RPE/choroid wholemounts was determined by TUNEL staining using an In Situ Cell Death Detection Kit , TMR Red (Roche Diagnostics, Burgess Hill, UK) as previously described (*7*).

Human iPSC derived RPE cell cultures were fixed with 2% PFA and permeabilization with 0.1% Triton X-100. After blocking with 10% normal goat serum, cells were incubated with a polyclonal rabbit anti-ZO-1 (1:200) overnight at 4°C, followed by labelling with goat anti-rabbit conjugated with Alexa Fluor 488 (1:400).

To stain mitochondria in frozen mouse retina sections, PFA-fixed samples were washed in PBS and incubated with 100 nM of MitoView Green (Insight Biotechnology, Wembley, UK) for 30 minutes at RT. After wash and counterstaining with DAPI, the samples were mounted for observation under confocal microscopy.

#### *Western blot*

Protein extraction from tissue or cells was prepared using Pierce® RIPA lysis buffer containing Halt™ Protease and Phosphatase Inhibitors (Thermo Fisher Scientific). Protein concentrations were measured using Pierce™ BCA protein assay kit (Thermo Fisher Scientific). 5-15 µg protein for each sample was mixed with Tris-Glycine SDS Sample Buffer (1:2) and reducing reagent (1:10, Thermo Fisher Scientific), followed by denaturing at 80°C for 2 min. After separation using Novex™ 4-20% Tris-Glycine Mini Gels (Thermo Fisher Scientific), proteins were transferred to PVDF membrane (Thermo Fisher Scientific), before blocking with 5% w/v milk in Tris-Buffered Saline (TBS) + Tween 20 (TBS-T; 0.1% v/v). Blots were incubated with primary antibodies for IRAK-M (1:2000), c-Jun (1:1000), phosphor-c-Jun (1:1000), c-Fos (1:1000), phosphor-c-Fos (1:1000), p21 (1:1000), lamin B1 (1:1000) or β-actin (1:2000) at 4 °C overnight. After thorough washing, blots were incubated with appropriate secondary antibody, anti-rabbit HRP (1:2000) or anti-mouse HRP (1:2000; both from Cell Signaling Technologies, London, UK). Chemiluminescence was detected by Amersham ECL reagents (Sigma-Aldrich) and developed using Hyperfilm™ ECL film (Sigma-Aldrich) and X developer.

#### *Multiplex cytokine array and enzyme-linked immunosorbent assay (EIA)*

Supernatants of ARPE-19 or BMMΦ cultures, or mouse sera prepared from lateral tail vein sampling were examined for the concentration of inflammatory cytokines using the LEGENDplex Human or Mouse Cytokine Array Kit (BioLegend, London, UK) according to the manufacturer's instructions. The concentration of HMGB1 in ARPE-19 cell culture supernatant was determined by a direct EIA using a polyclonal anti-HMGB1 (Abcam) as the manufacturer's protocol.

#### *Seahorse metabolic assay*

Seahorse XFp or XFe96 cell culture miniplates, sensor cartridges with utility plates, and all reagents for Mito Stress tests were obtained from Agilent Technologies. Different RPE cells were incubated in Seahorse XF DMEM (pH 7.4) containing 25 mM glucose, 1 mM pyruvate, and 2 mM glutamine in 37°C incubator without $CO_2$ for 45 min (*5*). Oligomycin (ATPase inhibitor, 1 μM), FCCP (protonophoric uncoupler, 0.5 μM) and antimycin A/rotenone (electron transport inhibitors, 1 μM) were injected where indicated and the oxygen consumption rate (OCR, pmol $O_2$/min) and extracellular acidification rate (ECAR, mpH/min) were measured in real time. The measurement rates were normalized by total protein content analyzed using a BCA assay. Metabolic parameters were calculated using the following formulae: nonmitochondrial respiration (minimum OCR after antimycin A/rotenone injection), basal respiration (difference between OCR before oligomycin and nonmitochondrial respiration), maximal respiration (difference between maximum OCR after FCCP injection and nonmitochondrial respiration), $H^+$ (proton) leak (difference between minimum OCR after oligomycin injection and nonmitochondrial respiration), ATP production (difference between OCR before oligomycin injection and minimum OCR after Oligomycin), spare respiratory capacity (difference between maximal respiration and basal respiration), glycolysis (maximum EACR before oligomycin injection), maximal glycolytic capacity (maximum ECAR after oligomycin injection) and glycolytic reserve (difference between maximal glycolytic capacity and glycolysis) (*5*).

#### *Senescence associated β-galactosidase staining*

Primary murine RPE cells cultured in laminin-precoated 8-well chamber slides (Corning GmbH, Wiesbaden, Germany) were treated with 0.25 mM PQ or $H_2O_2$ for 2 hours each day for a total of 7 days for induction of senescence (*6*). A fluorescence-based live cell senescence β-galactosidase (SA-β-gal) assay kit (Enzo Life Sciences, Exeter, UK) was used to quantify cellular senescence according to the manufacturer's instructions. Briefly, the cells were incubated with Pretreatment Solution at 37$^oC$ for 2 hours, followed by addition of SA-β-gal Substrate Solution (1:200). After a further 4 hours incubation, the cells were thoroughly rinsed with PBS and captured via the GFP channel (480 nm excitation, 520 nm emission) on an Evos FL Fluorescence Microscope.

#### *Chromatin immunoprecipitation (ChIP)*

A One-step ChIP kit (Abcam) was used to identify whether AP-1 subunits c-Jun and c-Fos are transcription factors of IRAK-M in RPE cells. Subconfluent ARPE-19 cells were fixed with 1% formaldehyde for 15 min and quenched with 0.125 M glycine. Chromatin was isolated by adding chromatin lysis buffer, followed by disruption with a Dounce homogenizer. Lysates were sonicated to shear the DNA to an average length of 300–500 bp. Cross-linked chromatin fragments (from $1 \times 10^6$ cells) were immunoprecipitated with ChIP-grade antibodies against c-Jun or c-Fos (Cell Signaling, London, UK), or irrelevant IgG control. The precipitated DNA was amplified by PCR to amplify the IRAK-M Transcription Start Site (TSS). The primer sequences (*8*) are as follows: forward 5'-TGTGGCCAGGCGGACGCAG-3'; reverse: 5'-AGGTCGAACAGCAGCGTGT-3′.

#### *Cytotoxicity assay*

At various time points, RPE cell culture supernatants were collected and chemical-induced RPE cytotoxicity was assessed using a Lactate Dehydrogenase (LDH) detection kit (Abcam) according to manufacturer's instructions. Activity of released LDH was normalized to the LDH value of RPE lysates (100% cytotoxicity).

#### *Detection of autophagy flux*

The formation of autophagosome and autolysosome in RPE cells was monitored through LC3B protein localization using the Premo™ Autophagy Tandem sensor RFP-GFP-LC3B Kit (Thermo Fisher Scientific) according to the manufacturer's instructions. The RFP-GFP-LC3B sensor kit uses the high transduction efficiency and minimal toxicity of BacMam 2.0 expression technology, enabling the detection of LC3B positive, neutral pH autophagosomes in green fluorescence (GFP) and LC3B positive acidic pH autolysosome in red fluorescence (RFP). The RFP and the GFP genes included in this chimera are TagRFP and Emerald GFP, respectively. Briefly, ARPE-19 cells were transduced with a mixture of TagRFP-LC3B and Emerald GFP-LC3B at a MOI of 30 in cell culture medium overnight, before the addition of chemicals for 24 hours. LC3B-positive puncta (green for autophagosomes and red for autolysosomes) were analyzed using fluorescence microscopy and quantified using Image J.

#### *Quantitative RT-PCR (QRT-PCR)*

Total RNA was isolated using TRIzol reagent (Thermo Fisher Scientific). One µg of total RNA was treated with RQ1 RNase-free DNase before reverse-transcription using the ImProm-II™ Reverse Transcription System (Promega). cDNA was amplified using the PowerUp SYBR® Green PCR Master Mix Reagent (Thermo Fisher Scientific) on a QuantStudio Real-Time PCR System. Primer sequences were designed using the Primer-BLAST (NCBI): Mouse *Irak3*, forward 5'- GACCAGCTCCAACCCAAACT, reverse 5'- GCCACCGCCGGTCATATTTA; Human *Irak3*, forward 5'- CCCACTCCCTTGGCACATTC, reverse 5'-AGCATGGTTGAACGTTGTGC; Mouse *Irak1*, forward 5'-CAGAGGTGGAACAGCTATCAAG, reverse 5'-CATTGGGCAAGAAGCCATAAAC; Mouse *Irak4*, forward 5'-AAAGGACAGGACATCCGTAATG, reverse 5'-TCGCTGGACTCTACACTTCT; mouse *Rps29*, forward 5'-ACGGTCTGATCCGCAAATAC, reverse 5'-ATCCATTCAAGGTCGCTTAGTC.

#### *In vivo induction of retinal degeneration*

Retinal degeneration in WT or *Irak3*$^{-/-}$ mice were induced by paraquat (PQ, a prooxidant) or light-induced oxidative damage as previously described (*9, 10*). Mice aged 8 weeks were anesthetized by intraperitoneal injection of 200 µl of Vetelar (ketamine hydrochloride 100 mg/ml, Pfizer, Sandwich, UK) and Rompun (xylazine hydrochloride 20 mg/ml, Bayer, Newbury, UK) mixed with sterile water in the ratio 0·6:1:84. Pupils were dilated using 1% tropicamide and 2.5% phenylephrine (both from Chauvin, Essex, UK). A drop of Viscotears (Novartis, London, UK) was then applied to cover the surface of the eye before the following procedures.

For PQ-induced retinal degeneration (*9*), administration of PQ (0.375-1.5 mM, Sigma-Aldrich, Poole, UK) or PBS in contralateral eye, was delivered by 2 µl intravitreal injection conducted under a surgical microscope.

For light-induced retinal degeneration (LIRD) (*10*), fundus camera-delivered intense light was delivered to retinas through a Nikon D80 digital camera that was connected to an endoscope with a 5-cm long teleotoscope. The position of the mouse was adjusted using stage controls to allow the cornea to contact the end of teleotoscope and position the optic disk at the centre of the fundus image. Light was applied to one eye at an intensity of 100 klux for a one-time exposure of 20 minutes. The light intensity was regularly measured using a light meter to ensure equal illumination. The contralateral eye was left without light challenge as control.

***Optical coherence tomography (OCT)***

At selected time-points during lifespan or after induction of retinal degeneration in mice, pupils were dilated, and animals anaesthetised by isofane for clinical assessment. The Micron IV retinal imaging microscope (Phoenix Research Laboratories, Pleasanton, CA) was used to capture OCT scans, and brightfield and fluorescence fundal images. Prior to imaging, the Micron IV CCD and OCT were calibrated in accordance with the manufacturer's protocol. The gain was set to +3 dB and the FPS to 15, or +12 dB and 2 for brightfield and GFP fluorescence imaging, respectively. OCT images were used to evaluate the retinal structure and thickness using ImageJ.

***Plasmid and siRNA transfection***

To activate IRAK-M or c-Jun expression in ARPE-19 cells, the cells were plated in 48-well plates for reach 70-80% confluence. The CRISPR activation plasmid (Santa Cruz Biotechnology) was used to upregulate the expression of endogenous gene expression. For each transfection, 0.3 µg of plasmid DNA were mixed with 1.5 µl Lipofectamine 3000 and 1 µl P3000 reagent and left for 15 min. The transfection complex was then added to the cells and left for 48 hours with reduced serum (1% FBS), followed by western blot analysis of protein expression. Non-targeting CRISPR Plasmid (Santa Cruz Biotechnology) serves as a negative control.

To induce stable expression of exogenous IRAK-M gene in B6-RPE07 cells, 70% confluent cells in 48-well plates were transfected with 0.3 µg of pUNO1 bearing the human or mouse IRAK-M gene (InvivoGen, Toulouse, France) using Lipofectamine 3000 as above. Two days post transfection, the media were replaced by selective culture media containing 10 µg/ml Blasticidin. The stable transfectants were selected and expanded over 3 weeks, and stable IRAK-M expression were determined by qRT-PCR or western blot.

To knockdown IRAK-M expression in ARPE-19 cells, the siRNA specific to human IRAK-M (Santa Cruz Biotechnology, Heidelberg, Germany) was utilized according to the manufacturer's instructions. The siRNAs were mixed with Lipofectamine 3000 reagent (Thermo Fisher Scientific) to form the transfection complex, prior to addition to RPE culture medium at a final concentration of 40 nM. Non-silencing siRNA was used as a negative control. IRAK-M expression was determined by western blot 48 hours post transfection.

#### *Subretinal administration of AAV*

Plasmid vectors AAV.CMV.hIRAK3.WPRE and control vector AAV.CMV.ORF-stuffer.WPRE were designed using VectorBuilder design software and subsequent cloning and AAV2 vectors were produced by VectorBuilder Inc. Viral titres were achieved by qPCR of the ITR sequence. Male C57BL/6J mice aged 8 weeks or *Irak3*$^{-/-}$ mice aged 2-4 months were anesthetized using an intraperitoneal injection of Vetelar/Rompun. Pupils were dilated using 1% tropicamide and 2.5% phenylephrine (Chauvin, Essex, UK). A drop of Viscotears was then applied to cover the surface of the eye before AAV administration. The trans-scleral subretinal injections were 2 μL in volume at the titers indicated and were delivered using an operating microscope and a 33G needle on a microsyringe under direct visualization (Hamilton Company, Reno, NV, USA). 1% Chloramphenicol ointment (Martindale Pharma, Wooburn Green, UK) was applied topically following injection.

### Supplementary Table

Table S1. Complete list of non-synonymous variants in the IRAK3 gene region extracted from the IAMDGC data (*1*).

| SNP | CHR | POS | MAF_IAMDGC | CONSEQUENCE |
|---|---|---|---|---|
| rs11465972 | 12 | 66,622,071 | 0.0001 | missense_variant |
| rs1168773 | 12 | 66,590,732 | 0.0056 | missense_variant |
| rs137909830 | 12 | 66,597,646 | 0.0006 | stop_gained |
| rs138274217 | 12 | 66,641,863 | <0.0001 | missense_variant |
| rs138984535 | 12 | 66,638,363 | 0.0015 | missense_variant |
| rs139342884 | 12 | 66,610,970 | 0.0005 | missense_variant |
| rs140116682 | 12 | 66,641,876 | <0.0001 | stop_gained |
| rs140228894 | 12 | 66,622,105 | <0.0001 | missense_variant |
| rs140671957 | 12 | 66,639,014 | 0.0001 | missense_variant |
| rs142201371 | 12 | 66,590,660 | <0.0001 | missense_variant,splice_region_variant |
| rs143857306 | 12 | 66,638,962 | 0.0001 | missense_variant |
| rs144713310 | 12 | 66,638,878 | 0.0001 | missense_variant,splice_region_variant |
| rs145220063 | 12 | 66,597,644 | 0.0013 | missense_variant |
| rs146096735 | 12 | 66,641,657 | <0.0001 | missense_variant |
| rs146120640 | 12 | 66,638,926 | 0.0006 | missense_variant |
| rs148849112 | 12 | 66,641,523 | <0.0001 | missense_variant |
| rs150324935 | 12 | 66,620,495 | <0.0001 | splice_polypyrimidine_tract_variant,splice_region_variant,intron_variant |
| rs188575230 | 12 | 66,639,011 | 0.0002 | missense_variant |
| rs199538395 | 12 | 66,639,013 | 0.0001 | missense_variant |
| rs200460084 | 12 | 66,638,999 | 0.0001 | missense_variant |
| rs200549011 | 12 | 66,605,332 | <0.0001 | missense_variant |
| rs200655368 | 12 | 66,641,908 | <0.0001 | missense_variant |
| rs201109976 | 12 | 66,597,523 | 0.0002 | missense_variant |
| rs34272472 | 12 | 66,638,879 | <0.0001 | missense_variant,splice_region_variant |
| rs34443407 | 12 | 66,597,607 | 0.0003 | missense_variant |
| rs34682166 | 12 | 66,605,300 | <0.0001 | missense_variant |
| rs35823766 | 12 | 66,622,068 | 0.0001 | missense_variant |
| rs76408141 | 12 | 66,605,329 | <0.0001 | stop_gained |

The 18 variants used in the gene burden test are shown in black text, while the variants not detected in both AMD cases and controls are shown in grey text.

| | |
|---|---|
| SNP | Rs-identifier on GRCh37 |
| CHR | Chromosome on GRCh37 |
| POS | Position on GRCh37 |
| MAF_IAMDGC | Allele frequency of the minor allele in the IAMDGC imputed data set |
| CONSEQUENCE | Functional consequence: either missense or stop gained indicating a nonsynonymous variant |

### Supplementary Figures

**Fig. S1**

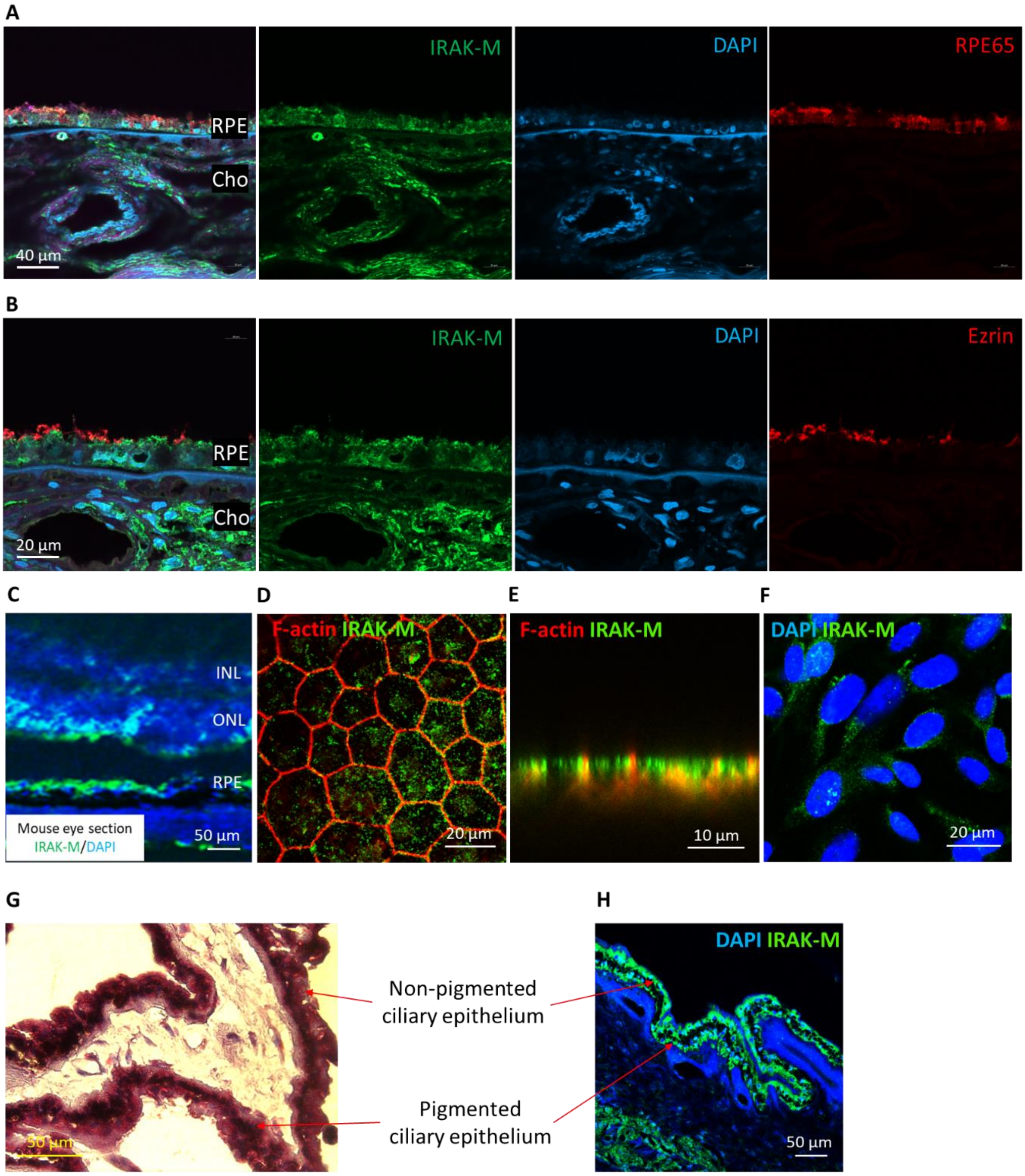


**Fig. S1. Immunostaining of IRAK-M in human and mouse eye tissues and ARPE-19 cells.** (**A&B**) Human RPE/choroidal cryosections from a 73-year-old male donor (no recorded ocular disease, exenterated eye fixed at time of surgery) were fixed in 4% PFA and lipofuscin autofluorescence was blocked by Sudan black B. The specimens were then stained for IRAK-M and RPE65 (**A**) and Ezrin (marker for RPE microvilli, **B**), counter-stained with DAPI, and observed under confocal microscopy. (**C**) Representative immunofluorescence image of retinal sections of adult C57BL/6J mice stained for IRAK-M. (**D**) Confocal image of mouse RPE/choroidal flatmount stained for IRAK-M and F-actin, and (**E**) side view of Z-stacks of the image. (**F**) Confocal image of ARPE-19 cells stained for IRAK-M. (**G**) ABC-AP IHC and (**H**) immunofluorescence images of paraffin-embedded human eye sections from a non-AMD donor (59-year old) show IRAK-M immunopositivity in both non-pigmented and pigmented ciliary epithelium. The ciliary pigmented epithelium has no autofluorescence.

Fig. S2

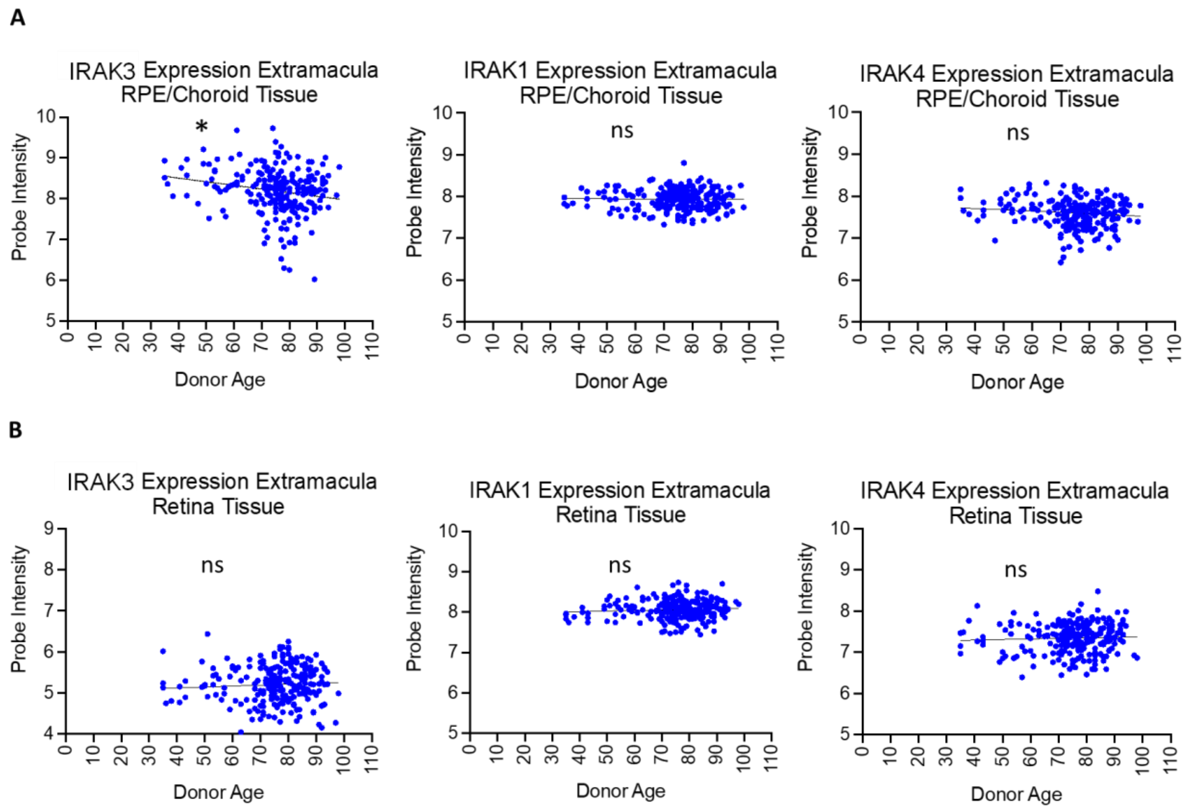


**Fig. S2. Correlation analysis of transcriptome of IRAKs in human extramacula RPE/choroid and retina with age.** Affymetrix chip-based transcriptome analyses show an age-dependent reduction in the expression level of IRAK3 mRNA in extramacular RPE/choroid tissues (**A**), but not in the retina (**B**). IRAK1 or IRAK4 mRNA level remains no change during ageing in RPE/choroid (**A**) and retina (**B**).

Fig. S3

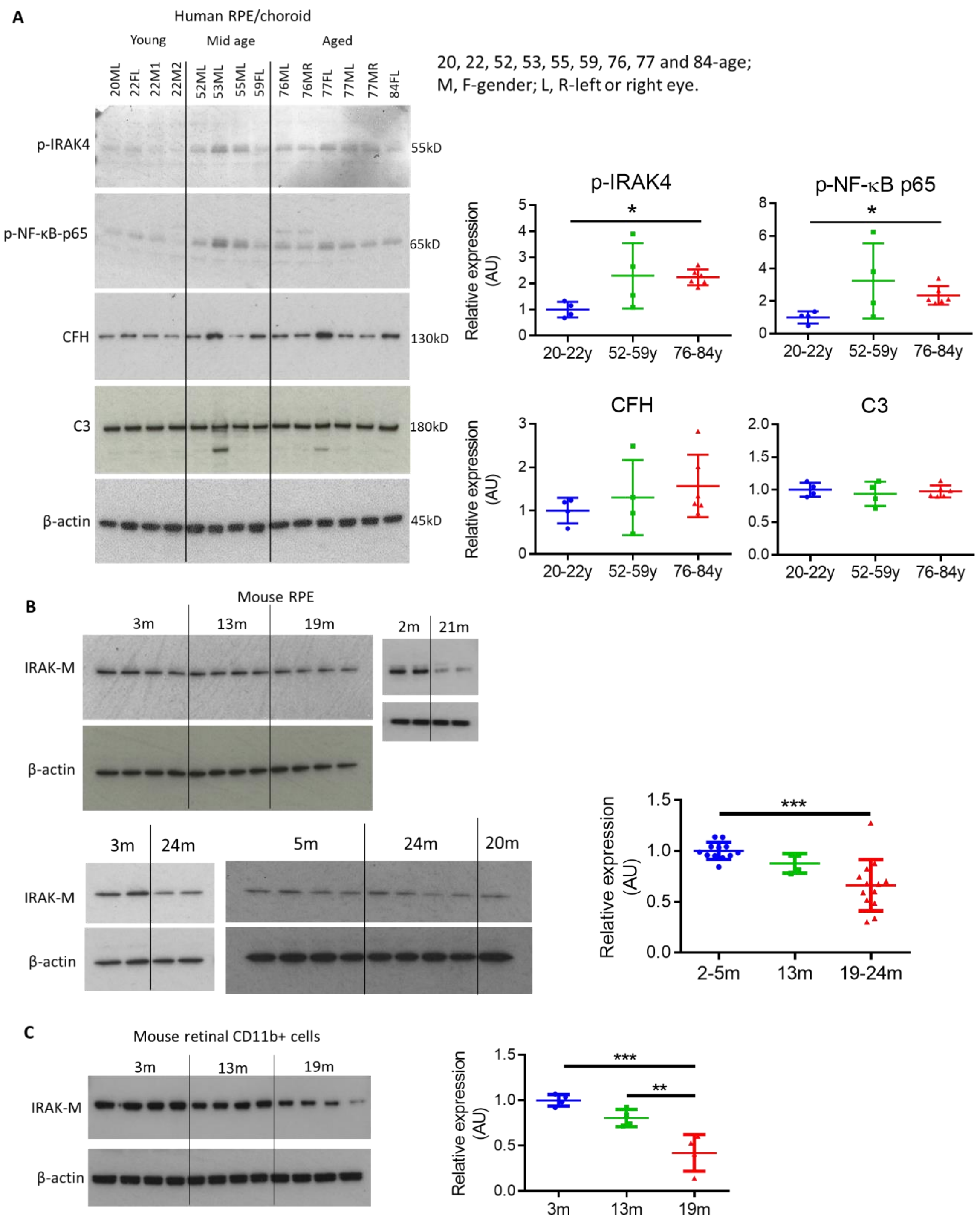


**Fig. S3. Western blots and densitometry analyses for age-associated changes in human and mouse ocular samples.** (**A**) Comparison of expression levels of phospho-IRAK4, phospho-NF-κB-p65, C3 and CFH in different age groups of human RPE/choroidal lysates (no recorded eye disease (n=4 or 6). (**B**) Comparison of IRAK-M expression levels in different age groups of RPE lysates from WT C57BL/6J mice (n=4-13). (**C**) Comparison of IRAK-M expression levels in different age groups of retinal CD11b+ cell lysates from WT C57BL/6J mice (n=4). *P < 0.05; **P < 0.01; ***P < 0.001. Comparison by one-way ANOVA.

Fig. S4

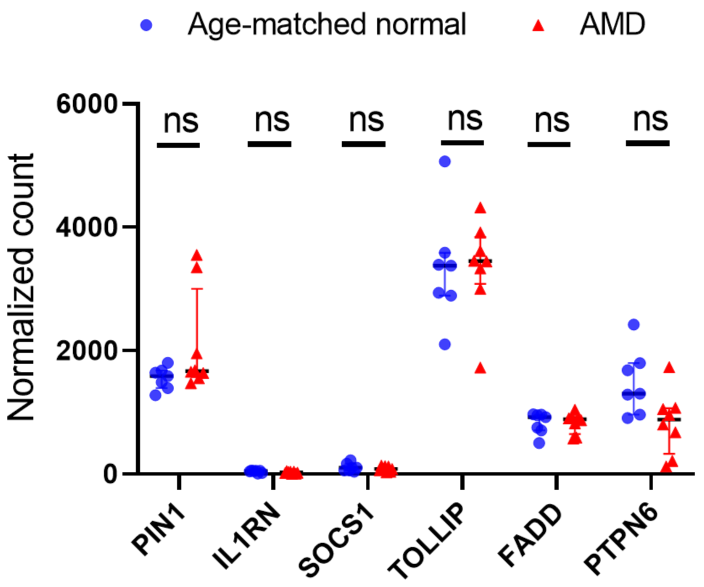


**Fig. S4. Comparison of PORT-normalized mRNA expression of known negative regulators for TLR/IL-1R/MyD88/IRAK1/4 signalling in RPE/choroid/sclera specimens between AMD and age-matched normal controls.** RNA-Seq data (GSE99248) analysis shows that there are no significant differences of PIN1, IL1RN, SOCS1, TOLLIP, FADD, and PTPN6 between AMD and controls. ns, nonsignificant. Comparison by two-way ANOVA (n=7-8).

Fig. S5

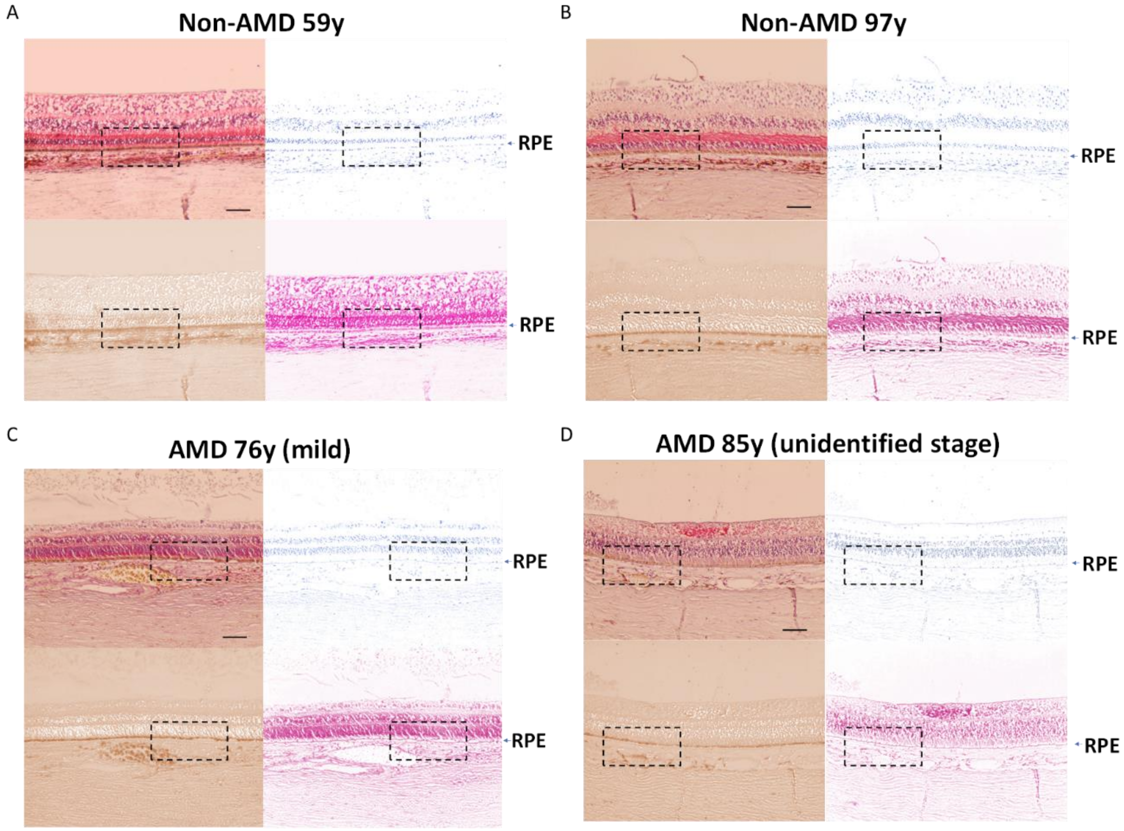


**Fig. S5. Representative IHC images show IRAK-M immunopositivity using paraffin-embedded human eye sections.** Slides were stained for IRAK-M, visualized using ABC-AP method, and counterstained with Hematoxylin. The original IHC images were color-deconvoluted using Fiji to separate IRAK-M staining (red), pigment (brown) and nuclei (blue). The images are from two non-AMD retinas from a 59-year old female donor (**A**) and a 97-year old female donor (**B**), respectively, and a mild AMD retina from a 76-year old male donor (**C**) and an unidentified stage AMD retina from an 85-year old male donor (**D**). Scale bars = 100 µm. Magnified boxed regions of IHC images (**A**-**D**) are shown in main Fig. 2B.

Fig. S6

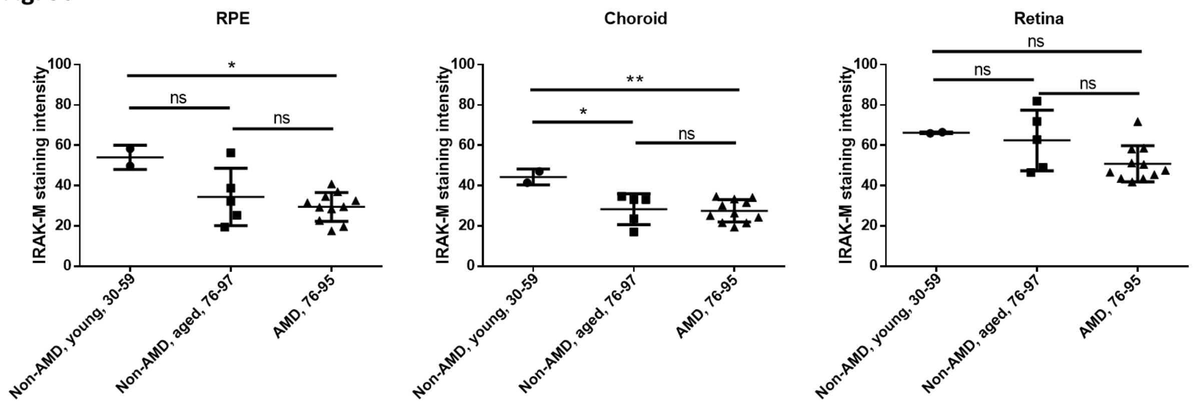


**Fig. S6. Comparison of IRAK-M expression at the extramacular areas of paraffin-embedded human eye sections.** Quantification of mean IRAK-M staining intensities of RPE, choroid and retina at the extramacular areas is shown (n=2 for young controls, n=5 for old controls and n=11 for AMD). *P < 0.05; **P < 0.01; ns, nonsignificant. Comparison by one-way ANOVA.

Fig. S7

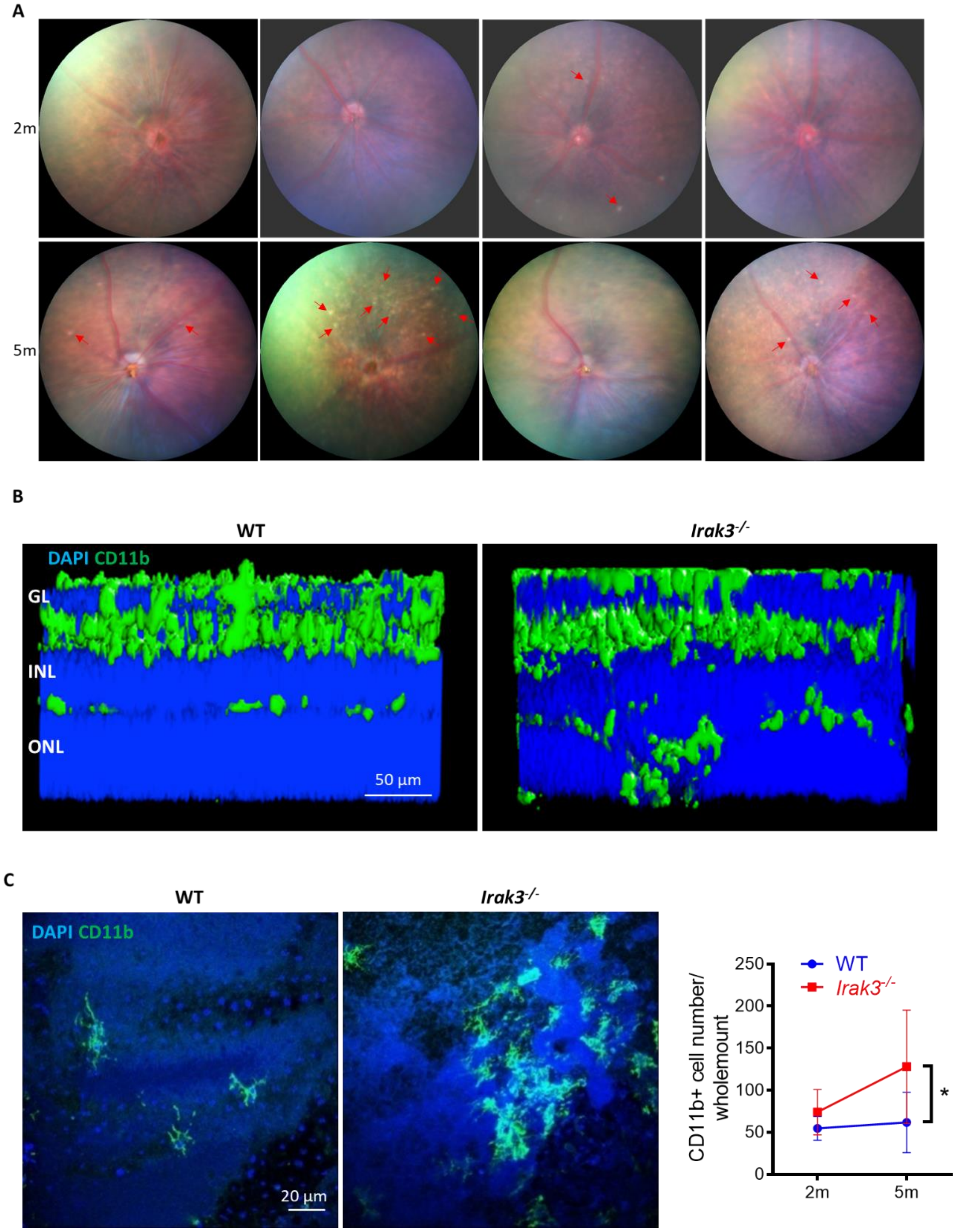


**Fig. S7. *Irak3*$^{-/-}$ mice exhibit abnormal retinal spots and myeloid cell response.** (**A**) Representative fundal images show increased incidence of retinas presenting scatted white spots (red line arrow) in *Irak*$^{-/-}$ mice aged 5m, which is not evident at 2m. (**B**) Z-stacks of confocal images of retinal flatmounts demonstrate presence of CD11b+ myeloid cell populations in the outer retina in 5m-old *Irak3*$^{-/-}$ mice compared to WT counterparts. (**C**) Subretinal accumulation of CD11b+ myeloid cells was assessed by immunostaining on RPE/choroidal flatmounts of *Irak3*$^{-/-}$ vs. WT mice (n=3-10). *P < 0.05 by two-way ANOVA.

Fig. S8

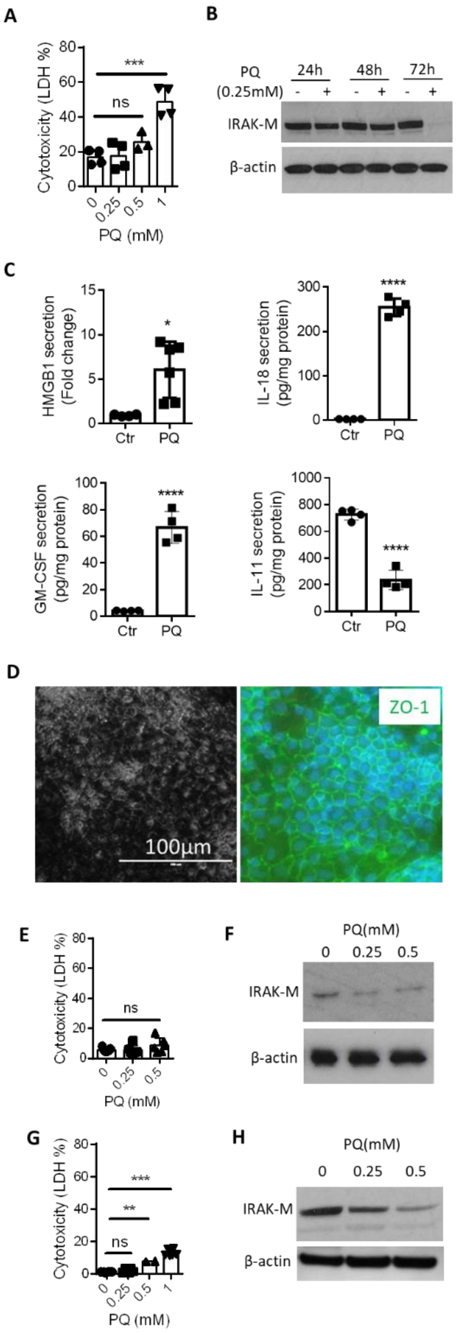


**Fig. S8.** ***In vitro*** **oxidative treatment downregulates IRAK-M expression in RPE cells.** (**A**) Human ARPE-19 cells were treated with various concentrations of a prooxidant, paraquat (PQ), for up to 72h. LDH release demonstrate dose-dependent cytotoxic effect of PQ after 72h (n=3-4). (**B**) Treatment of ARPE-19 cells with a subtoxic dose of PQ (0.25mM) reduces IRAK-M expression after 72h (Western blot), which is accompanied by (**C**) increased secretion of pro-inflammatory cytokines HMGB1 (EIA, n=4-6), IL-18 and GM-CSF, and decreased anti-inflammatory cytokine IL-11 (multiplex cytokine array, n=4). (**D**) Pigmented human iPSC-derived RPE cells with typical hexagonal morphology (ZO-1 stain). (**E**) LDH cytotoxicity assay determines sub-toxic doses of PQ on human iPSC-derived RPE after 72h treatment (n=5). (**F**) Western blot shows downregulated IRAK-M expression by 72h treatment of subtoxic PQ doses (0.25-0.5mM). LDH assay(**G**, n=3-6) and western blot (**H**) show curtailed IRAK-M expression in human primary RPE cells by 72h treatment of subtoxic PQ dose (0.25mM). *P < 0.05; **P < 0.01; ***P < 0.001; ****P < 0.0001; ns, nonsignificant. Comparison by one-way ANOVA (A, E and G) or unpaired two-tailed Student's t-test (C).

**Fig. S9**

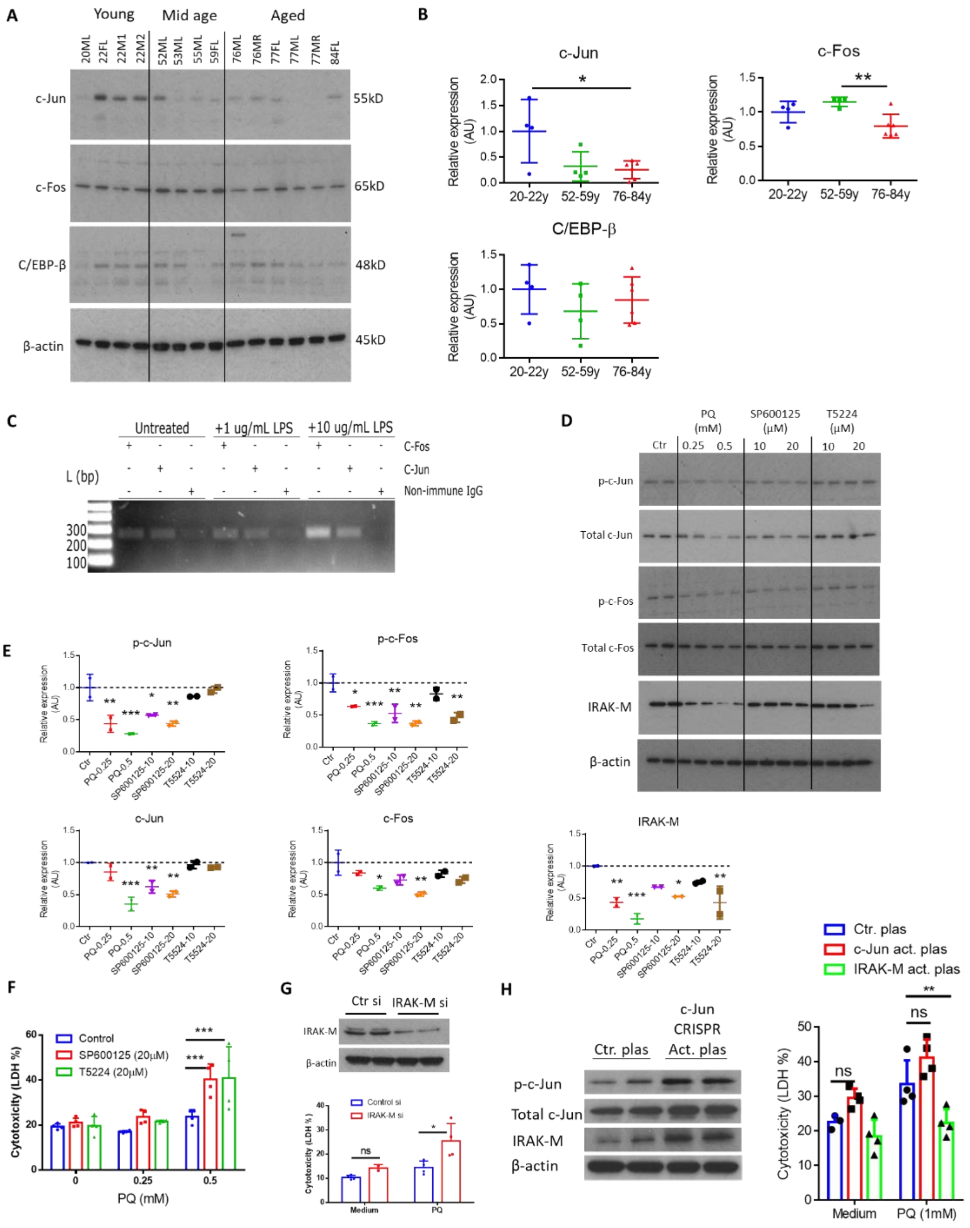


**Fig. S9. IRAK-M expression is regulated by AP-1 in RPE.** (**A**) Western blot and (**B**) densitometry analyses compare expression levels of c-Jun, c-Fos and C/EBP-β in RPE/choroidal lysates from human donor eyes with different ages (n=4-6). (**C**) ChIP assay of ARPE-19 cells shows the binding of AP-1 subunits c-Jun and c-Fos to IRAK-M promoter under resting condition, and at a higher extent, by LPS stimulation for 24h. Non-immune IgG was used as a negative control. (**D**) Western bolt and (**E**) densitometry analyses show PQ dose-dependent downregulation in phosphorylation of c-Jun and c-Fos, and total c-Jun and c-Fos expression, in parallel with downregulated IRAK-M expression in ARPE-19 cells. AP-1 subunit inhibitors SP600125 (primarily targeting c-Jun) or T5224 (for c-Fos) at 20 μM decrease IRAK-M expression. (**F**) LDH assay shows increased susceptibility of ARPE-19 cells in response to PQ treatment when c-Jun or c-Fos is inhibited (n=4). (**G**) IRAK-M knockdown (western blot) by siRNA exacerbates PQ- induced ARPE-19 cytotoxicity (n=4). (**H**) Overexpression of c-Jun by CRISPR/Cas9 activation plasmid induces total and phosphorylated c-Jun, and IRAK-M expression in ARPE-19 (Western blot). However, c-Jun overexpression does not prevent from PQ-induced cytotoxicity, in contrast to a protecting effect of IRAK-M overexpression via CRISPR/Cas9 (n=3-4). $^{*}P < 0.05$; $^{**}P < 0.01$; $^{***}P < 0.001$; ns, nonsignificant. Comparison by one-way ANOVA (B and E) or two-way ANOVA (F, G and H).

**Fig. S10**

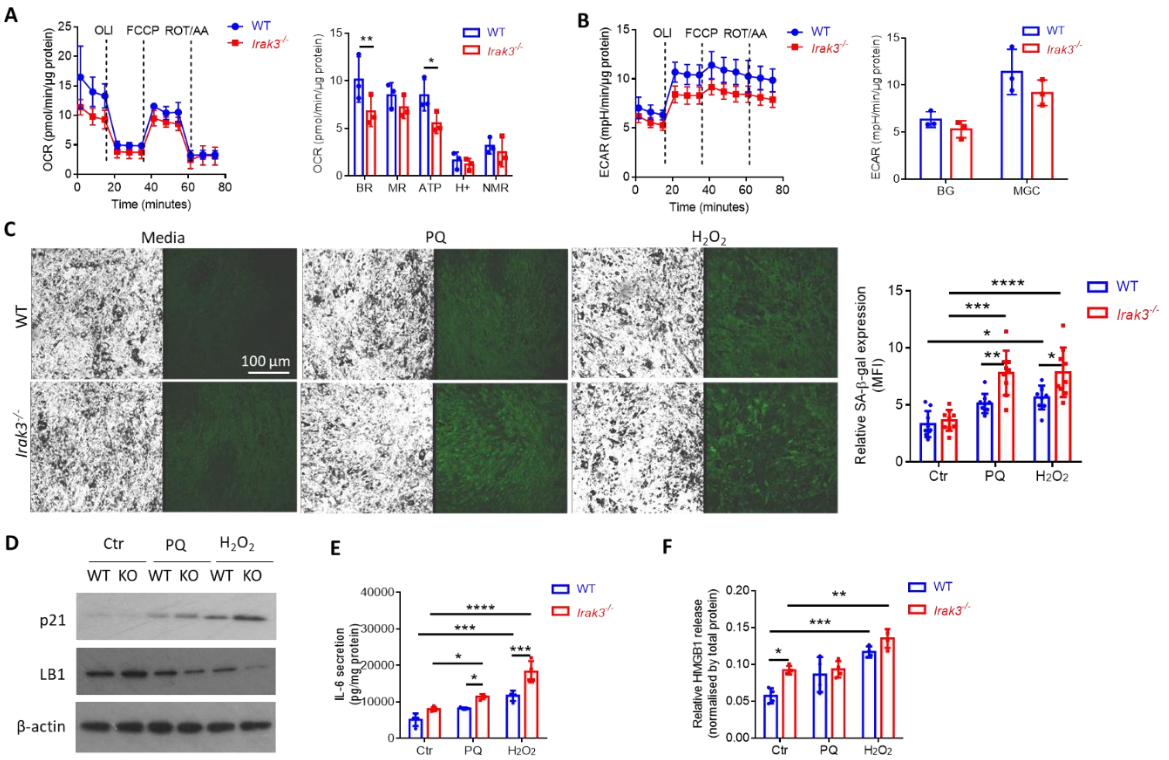


**Fig. S10. Absence of IRAK-M leads to impaired RPE cell homeostasis.** Primary RPE cells isolated from WT or *Irak*$^{-/-}$ mice (5m old) were subjected to Mitochondrial Stress Test using a Seahorse XFp Analyzer, and metabolic parameters calculated from OCR (**A**) and ECAR (**B**) profiles demonstrate decreased mitochondrial basal respiration (BR) and ATP production in *Irak*$^{-/-}$-RPE cells, despite no significant differences in maximal respiration (MR), proton leak (H+), non-mitochondrial respiration (NMR), basal glycolysis (BG) and maximal glycolytic capacity (MGC), between WT and *Irak*$^{-/-}$-RPE (n=3). (**C**) Mouse primary RPE cells subjected to a pulsed PQ or $H_2O_2$ treatment (repeated 2h treatment per day for a total of 7 days) were analyzed for cellular senescence using a fluorescence-based SA-β-gal assay. Mean fluorescence intensity quantified using Fiji demonstrates induced SA-β-gal signal in *Irak*$^{-/-}$ RPE cells (n=9). Oxidative stress-induced, and *Irak*$^{-/-}$-promoted RPE senescence was also confirmed by increased p21 and decreased Lamin-B1 expression (**D**), and enhanced secretion of proinflammatory cytokines IL-6 (**E**, n=4) and HMGB1 (**F**, n=4). *P < 0.05; **P < 0.01; ***P < 0.001; ****P < 0.0001. Comparison by two-way ANOVA.

**Fig. S11**

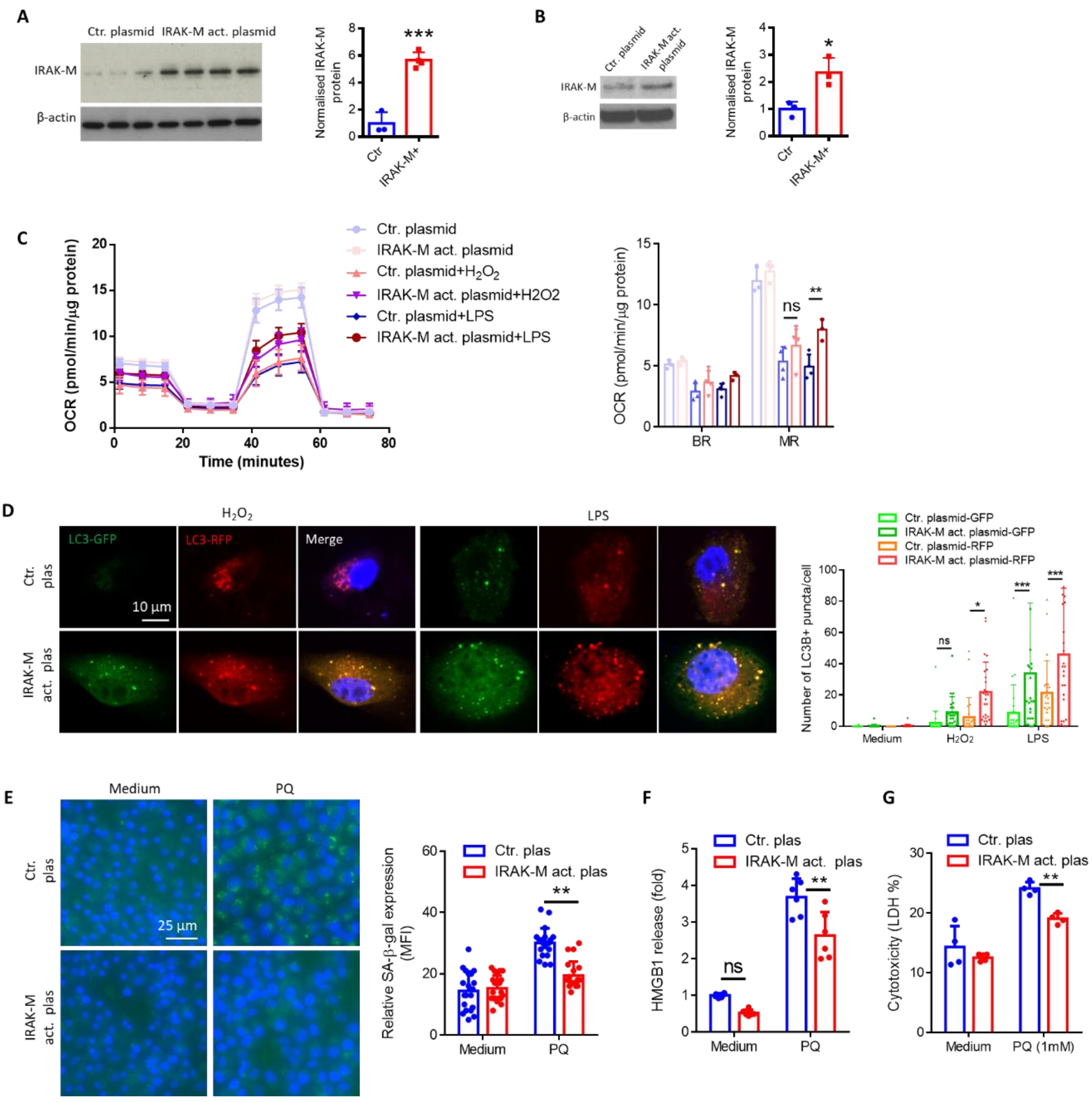


**Fig. S11. Overexpression of IRAK-M in RPE cells maintains cell homeostasis against stressors.** (**A**) Western blot analysis of IRAK-M expression in human iPSC-RPE cells following transfection with CRISPR/Cas9 activation plasmid for 48h (n=3 or 4). (**B**) Western blot analysis of IRAK-M expression in ARPE-19 cells following transfection with CRISPR/Cas9 activation plasmid for 48h (n=3). (**C**) OCR profile and parameter analysis show a partial inhibition of LPS-caused reduction in maximal respiration (MR) by IRAK-M overexpression in ARPE-19 cells (n=3 or 4). (**D**) Autophagy Tandem LC3B-GFP-RFP sensor assay and confocal imaging demonstrate that IRAK-M overexpression in ARPE-19 cells enhances the formation of LC3B-autophagosome (green) and LC3B-autolysosome (red) in response to $H_2O_2$ or LPS treatment (n=20-25). (**E&F**) PQ-induced ARPE-19 senescence is inhibited by IRAK-M overexpression, evidenced by reduced SA-β-gal activity (n=20, **E**) and HMGB1 release (n=6, **F**). (**G**) LDH cytotoxicity assay shows that IRAK-M overexpression in ARPE-19 cells inhibits PQ-induced cytotoxicity (n=4). *P < 0.05; **P < 0.01; ***P < 0.001; ns, nonsignificant. Comparison by unpaired two-tailed Student's t-test (A and B), or two-way ANOVA (C-G).

**Fig. S12**

**A**

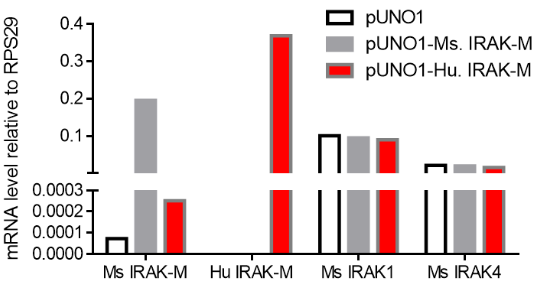


**B**

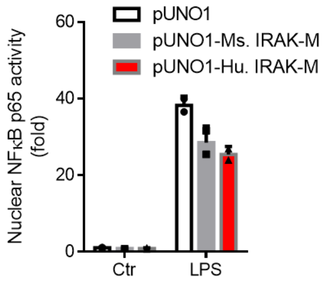


**Fig. S12. Overexpression of mouse or human IRAK-M in mouse RPE cells inhibits nuclear NF-κB activity.** Stably transfected cell colonies were selected from mouse B6-RPE07 cell line, and two new cell lines were established to persistently express mouse and human IRAK-M, respectively. Another new cell line bearing vehicle plasmid (pUNO1) was a transfection control. (**A**) qRT-PCR analysis shows overexpression of mouse or human IRAK-M gene in the cell lines, without changes in expression of IRAK1 or IRAK4. (**B**) The cell lines were stimulated with 1 µg/ml LPS for 30 min to activate TLR4/NF-κB signalling. NF-κB activity assay shows reduced nuclear NF-κB-p65 binding activity in cells overexpressing mouse or human IRAK-M (n=2).

**Fig. S13**

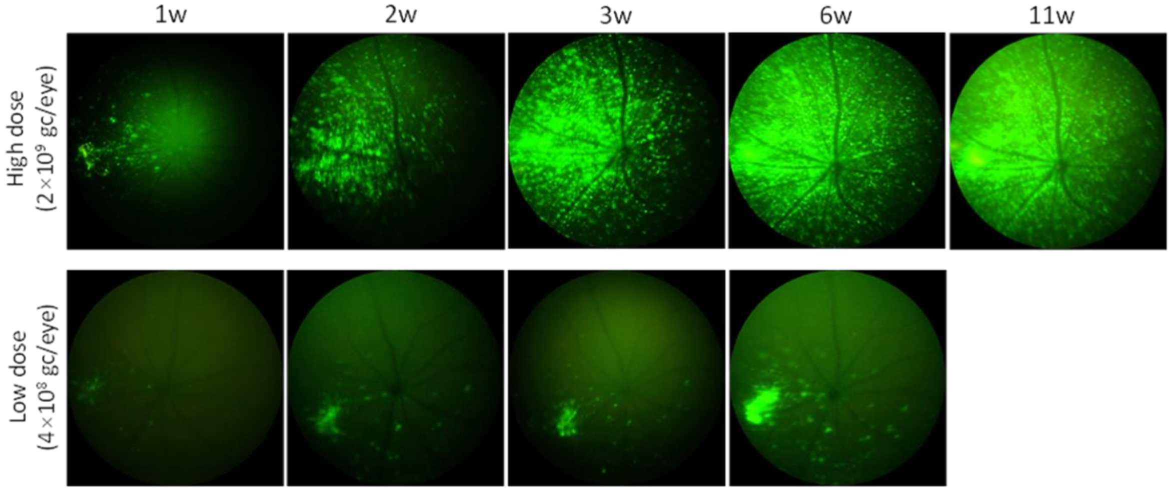


**Fig. S13. AAV2 dose-dependently transduces retinal tissues following subretinal administration.** A total of $2\times10^9$ (high dose) or $4\times10^8$ (low dose) genome copies (gc) of AAV2 encoding EGFP driven by the CMV promoter were injected into the subretinal space per eye in 8w-old WT mice. At 1-11 weeks post injection, viral dose-dependent retinal transduction was observed by fundal fluorescence imaging using a Phoenix MICRON® IV retinal imaging microscope.
